# Structural mechanism of TRPC3 inhibition by a potent and selective antagonist

**DOI:** 10.64898/2026.01.07.698177

**Authors:** Jodie Collingridge, Bethan A. Cole, John Liddle, Kelly M. Gatfield, Philip C. Biggin, Esther B. E. Becker

## Abstract

The transient receptor channel canonical family member 3 (TRPC3) is a non-selective, calcium-permeable cation channel within the larger TRP superfamily. TRPC3 is expressed in both excitable and non-excitable cells, where it integrates multiple signaling pathways and is directly activated by binding of diacylglycerol following activation of G-protein coupled receptors. Increased TRPC3 signaling is implicated in several diseases, highlighting TRPC3 as a potential therapeutic target. However, the development of TRPC3 inhibitors with suitable potency and selectivity has been challenging due to the channel’s high structural and sequence homology with related family members, particularly TRPC6. Here, we present a novel, selective and highly potent TRPC3 antagonist (GSK2820986A) featuring an anilino-thiazole pharmacophore. Through a combination of molecular dynamic simulations and functional assays, we identify a putative binding site for GSK2820986A in the S4-S5 pocket and propose a potential inhibitory mechanism. Our findings provide a powerful tool compound and mechanistic framework to advance the investigation and therapeutic targeting of TRPC3 in disease.

## Introduction

The transient receptor potential canonical family member 3 (TRPC3) belongs to the large, heterogeneous transient receptor potential (TRP) channel superfamily (1). TRPC3 is a tetrameric, non-selective, calcium-permeable cation channel that acts as a multifunctional signaling molecule downstream of G protein-coupled receptors (2, 3). The channel belongs to the TRPC family subgroup TRPC3/6/7, which share the highest sequence similarity at ∼70-80% (4). The TRPC3/6/7 subfamily is unique in that the primary mechanism of activation is thought to be direct binding of the lipid molecule diacylglycerol (DAG) in the fenestration between the S6 of one subunit and the pore-loop helix of another (5, 6). In addition, TRPC3 has also been shown to exhibit DAG-independent constitutive activity, more so than the other subfamily members (7, 8).

TRP channels have a highly conserved S1-6 transmembrane domain (TMD) architecture, which is broadly conserved across many other families within the larger P-loop channel superfamily to which the TRP family belongs (9) (Fig. 1). This architecture is characterized by the membrane-re-entrant pore-loop and either a voltage-sensing domain (VSD), in the case of voltage-gated P-loop family members, or an equivalent voltage-sensing-like domain (VSLD), in the case of TRP channels (9). TRPC3 contains these shared structures, as well as an intracellular domain arrangement that is common to the known TRPC structures (10). The intracellular domain contains multiple ankyrin repeat domains (ARD) and two characteristic C-terminal helices, ‘horizontal’ and ‘vertical’ (10). The C-terminal region of TRPC3 facilitates channel regulation via multiple mechanisms. These include inhibition through binding of calmodulin at the calmodulin/IP_3_R-binding (CIRB) domain, activation through IP_3_R binding at the CIRB, and inhibition through direct binding of calcium (11–13). Human TRPC3 is expressed throughout the body but has distinct regions of high expression and functional importance. TRPC3 is highly expressed in the nervous system, particularly in the cerebellum. Within the cerebellum, TRPC3 is strongly expressed in Purkinje cells, where the channel plays a key role in sensorimotor integration by mediating synaptic transmission and Purkinje cell activity (14, 15). Aberrant TRPC3 signaling is linked to cerebellar disease. In particular, gain-of-function (GOF) mutations in the gene encoding TRPC3 have been shown to cause cerebellar ataxia in the *Moonwalker (Mwk)* mouse model and spinocerebellar ataxia type 41 (SCA41) in humans (16, 17). The *Mwk* mutation (p.T561A) is located within the S4-S5 linker region of TRPC3, whereas the identified human TRPC3 variant (p.R677H) resides within the TRP helix (Fig. 1A). However, in 3D space, both mutations are close to each other and the TRPC3 pore (Fig. 1B) and both are thought to affect TRPC3 channel gating (2). Increased TRPC3 activity downstream of metabotropic mGluR1 receptor signaling also likely contributes to disease in other genetic subtypes of SCA (18, 19). Moreover, increased TRPC3 function has been linked to hyperexcitability and seizure activity in rodent models (20–22). Additionally, TRPC3 expression and activity in primary sensory neurons has been associated with chronic itch (23).Outside of the central nervous system, TRPC3 has been implicated in pathologies of the cardiovascular system including cardiac hypertrophy via activation of the calcineurin/nuclear factor of activated T-cells (NFAT) pathway (24), as well as cardiac fibrosis and atrial fibrillation (25, 26). TRPC3 may also contribute to the proliferation, differentiation and migration of various types of cancer cells (27–29).

**Figure 1.**
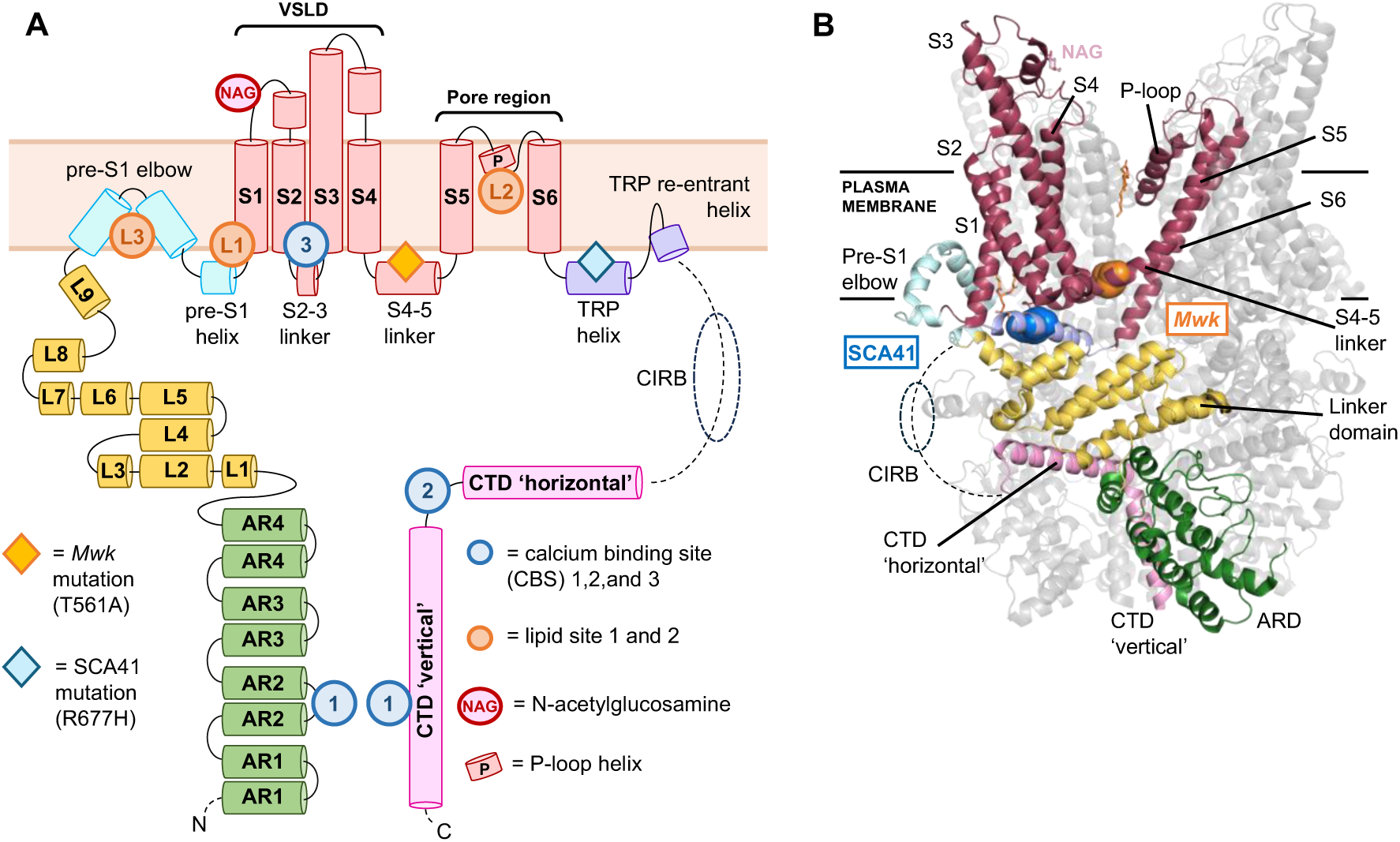
TRPC3 domain architecture and disease mutations. **(A)** Schematic of one TRPC3 subunit with sites of regulation and modification highlighted. AR, ankyrin repeat; CTD, C-terminal domain; CIRB, calmodulin/IP_3_R-binding domain; L, linker helix; NAG, N-acetylglucosamine; TRP, transient receptor potential domain; VSLD, voltage-sensing-like domain. The *Moonwalker (Mwk)* mutation (T561A) and SCA41 patient variant (R677H) both confer pathogenic gain of function to the channel. **(B)** TRPC3 subunit structure (PDB code 6CUD) with the plasma membrane indicated, labelled as in (A).

Given the association of TRPC3 with various pathologies, the development of TRPC3 functional modulators has been an area of intense interest. Although various TRPC3 channel agonists and antagonists have been published (2, 30, 31), their development has been hampered by the high sequence similarity between different TRPC channels, particularly between TRPC3 and TRPC6. Co-targeting of TRPC3 and TRPC6 channels is highly undesirable considering TRPC6’s role in a number of physiological processes including lung and kidney function and thus potential significant side effects (32, 33). One obstacle to the development of TRPC3-selective compounds is the incomplete understanding of TRPC3 structural and functional mechanisms and dynamics, in part due to the lack of an open-state structure for any of the TRPC3/6/7 family.

In this study, we present a highly potent and selective TRPC3 inhibitor, GSK2820986A (GSK-986), that arose from a drug discovery program based on aniline-thiazole pharmacophore-based blockers of TRPC3 and TRPC6 (34). We found that GSK-986 inhibited TRPC3 in the sub-nanomolar range and with ten-fold selectivity over TRPC6. We further identified the binding site of GSK-986 as the S4-S5 linker residue Q555. Molecular dynamics (MD) simulations revealed a potential mechanistic basis for the GSK-986-mediated TRPC3 inhibition via stabilization of the S4-S5 linker and thereby preventing transition to the active-state channel conformation. Together, our findings demonstrate that GSK-986 is a useful tool for selective TRPC3 inhibition and offer mechanistic insights that will advance development of selective TRPC3 antagonists for future clinical translation.

## Results

### GSK-986 is a potent and selective inhibitor of TRPC3

The GSK-986 compound (Fig. 2A) was developed as part of a series of anilino-thiazole pharmacophore compounds that showed TRPC3/TRPC6 inhibition with nanomolar potency and selectivity (34). Compounds were initially identified and optimized in a high-throughput screen using a carbachol-stimulated membrane potential FLIPR assay with human or rat TRPC3-overexpressing HEK293 cells. In this assay, GSK-986 was found to inhibit TRPC3 with pIC_50_ values of 8.15 (IC_50_=7nM, hTRPC3) and 8.7 (IC_50_=2 nM, rTRPC3), respectively (Fig. 2B). Moreover, the compound displayed about ten-fold selectively for TRPC3 over the related TRPC6 channel in the FLIPR assay (Fig. 2B). Inhibition of TRPC3 by GSK-986 was also evaluated using automated perforated patch clamp planar array electrophysiology (IonWorks Quattro) in human and mouse TRPC3-overexpressing HEK293 cells, following direct activation of TRPC3 with the previously described selective agonist GSK1702934A (35). These experiments confirmed that GSK-986 inhibits TRPC3 in the nanomolar range with pIC_50_ values of 7.43 (IC_50_=37 nM, hTRPC3) and 7.2 (IC_50_=63 nM, mTRPC3), respectively (Fig. 2B). Selectivity against TRPC6 was not assessed using this method. In additional experiments, GSK-986 showed low activity (IC_50_>10 μM, pIC_50_<5) against a panel of ion channels associated with cardiotoxicity (Fig. 2B).

**Figure 2.**
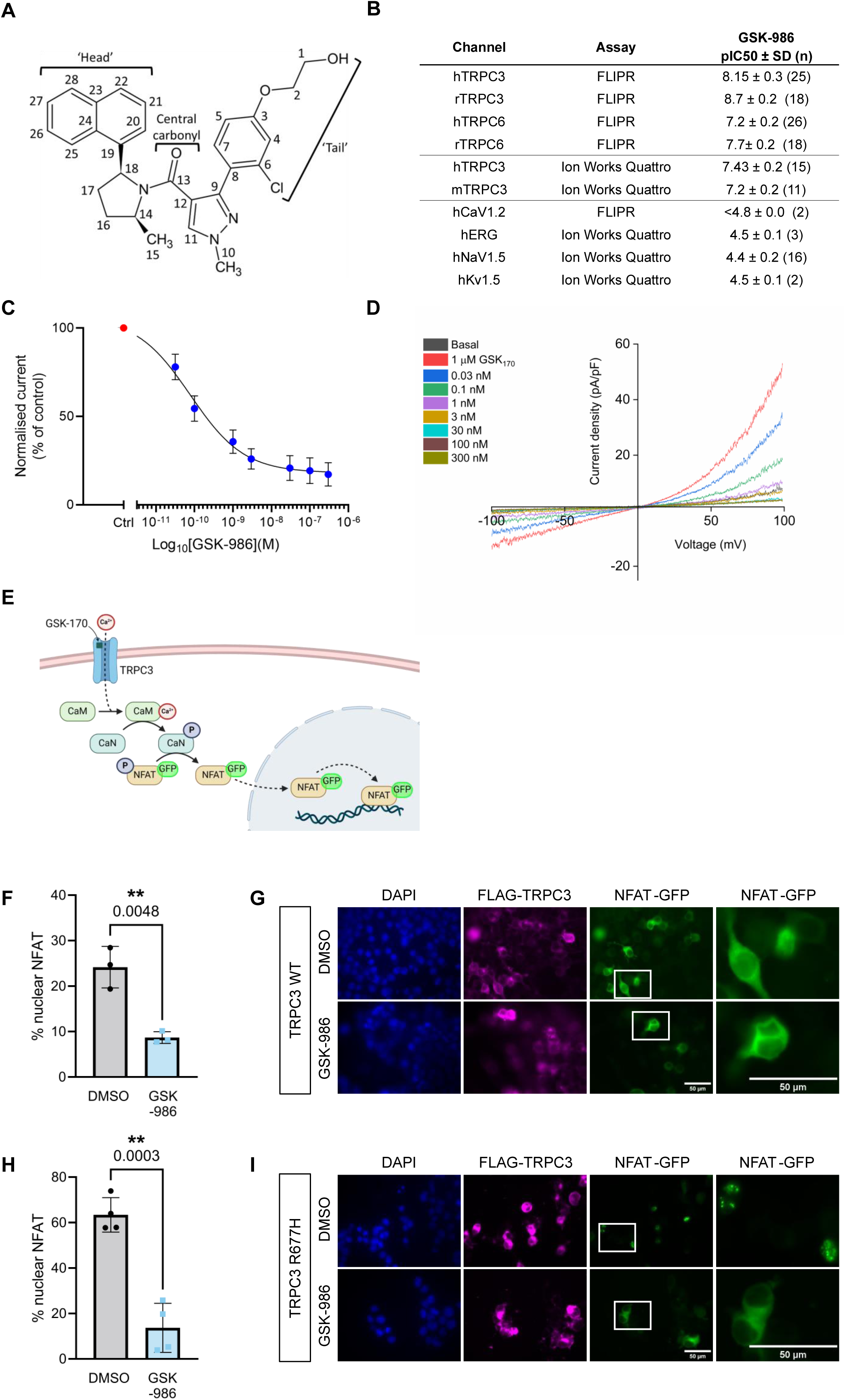
GSK-986 is a potent and selective inhibitor of TRPC3. **(A)** 2D chemical structure of TRPC3 inhibitor GSK2820986A (GSK-986). Drawing generated using ChemSketch. **(B)** pIC50 values (-logIC_50_) for GSK-986 obtained from high-throughput screening using FLIPR assay and automated planar perforated patch-clamp electrophysiology (Ion Works Quattro). **(C)** Dose-response curve for GSK-986 inhibition of TRPC3 expressed in HEK293T cells. Whole-cell patch-clamp recordings following activation with 1μM GSK1702934A in the presence of GSK-986. IC_50_=0.08 nM (n=5 cells). **(D)** Representative whole-cell currents recorded from HEK293T cells transiently transfected with TRPC3, activated using 1 μM GSK1702934A (GSK_170_) and exposed to 0.03-300 nM GSK-986 or DMSO control. **(E)** Schematic of the GFP-NFAT translocation assay. Upon activation of TRPC3 by GSK1702934A and subsequent increase of cytoplasmic calcium concentration, calcium binds and activates calmodulin (CaM), which phosphorylates and activates the phosphatase calcineurin (CaN). Phosphorylated CaN dephosphorylates GFP-tagged NFAT, which translocates to the nucleus. Figure created in Biorender. **(F)** GSK-986 significantly reduces nuclear GFP-NFAT localization in Neuro-2a cells transiently co-transfected with wildtype (WT) FLAG-TRPC3 and GFP-NFAT compared to DMSO vehicle-treated control. Cells were treated with 10 μM of the activator GSK1702934A and either 1 μM GSK-986 or an equivalent volume of DMSO. The percentage of nuclear-localized GFP-NFAT represents TRPC3 activity. Unpaired two-tailed T-test, **p<0.01, n=3 independent biological replicates, each being the percentage of cells with nuclear-located GFP-NFAT across 50-200 cells. Scatter points represent biological replicates and bar graph shows mean ± standard deviation (SD). **(G)** Representative images of NFAT-GFP translocation experiments quantified in (F). Treated cells were fixed and immunostained with antibodies against FLAG and GFP. Nuclei are visualized with DAPI in blue. Image windows are expanded to show NFAT localization in individual cells. Scale bars: 50 μm. **(H)** GSK-986 significantly reduces nuclear NFAT localization in Neuro-2a cells transiently transfected with TRPC3 harboring the SCA41 GOF disease mutation p.R677H. Cells were treated with either 1 μM GSK-986 or an equivalent volume of DMSO. The percentage of nuclear-localized GFP-NFAT represents TRPC3 activity. Unpaired two-tailed T-test, ***p<0.001, n=4 independent biological replicates, each being the percentage of cells with nuclear-located GFP-NFAT across 50-200 cells. Scatter points represent biological replicates and bar graph shows mean ± SD. **(I)** Representative images of NFAT-GFP translocation experiments quantified in (H). Treated cells were fixed and immunostained with antibodies against FLAG and GFP. Nuclei are visualized with DAPI in blue. Image windows are expanded to show NFAT localization in individual cells. Scale bars: 50 μm.

Subsequently, inhibition of hTRPC3 by GSK-986 was investigated in lower throughput cell-based assays. Using manual whole-cell patch-clamp electrophysiology in HEK293T cells transiently transfected with TRPC3, GSK-986 strongly inhibited TRPC3 currents with a pIC_50_ of 10.097 (IC_50_=0.08 nM) (Fig. 2C, D). To test the effect of GSK-986 inhibition on TRPC3-mediated calcium influx, we used a previously established cell-based NFAT-GFP reporter assay (17). Mouse neuroblastoma Neuro-2a cells were transiently co-transfected with FLAG-tagged hTRPC3 and GFP-tagged NFAT. Pharmacological activation of TRPC3 with the agonist GSK1702934A results in calcium influx and subsequent nuclear translocation of NFAT (Fig. 2E). Incubation with GSK-986 led to significant reduction in NFAT translocation following TRPC3 activation with GSK1702934A (Fig. 2F, G). We also tested the effect of the inhibitor on enhanced TRPC3 activity caused by the SCA41 GOF patient mutation. Overexpression of the constitutively active SCA41-associated TRPC3 mutant p.R677H resulted in strong nuclear translocation of GFP-NFAT as reported previously (17). The enhanced TRPC3 activity was significantly inhibited by incubation with GSK-986 (Fig. 2H, I). Together, these findings demonstrate that GSK-986 is a selective, direct, and potent inhibitor of TRPC3.

### Computational methods predict the vanilloid-like S4-S5 pocket as GSK-986 binding site

Based on the availability of TRPC3 structures obtained from cryo-electron microscopy (10, 36), we next sought to identify possible binding sites for GSK-986. To gain an idea of the stability of the TRPC3 and to relax any effects of crystal-lattice packing, we performed a 100-ns conventional molecular dynamics (MD) simulation using the 6CUD PDB structure (Uniprot Q13507-1) (10) (Figs. 3 and S1A). Docking of GSK-986 was then performed against the whole protein from the final structure of one MD run using QuickVina-W (36). Several possible binding sites were found, but ligands were found to cluster most frequently in two of them: the S4-S5 binding pocket and an extracellular site in the voltage-sensing domain (VSLD) (Fig. 3B). In three out of four subunits, ligands occupied the S4-S5 pocket, in comparable conformations (Fig. 3C). Some ligand poses were also found in the two lipid sites, L1 and L2, identified in the 6CUD cryo-EM structure (10).

**Figure 3.**
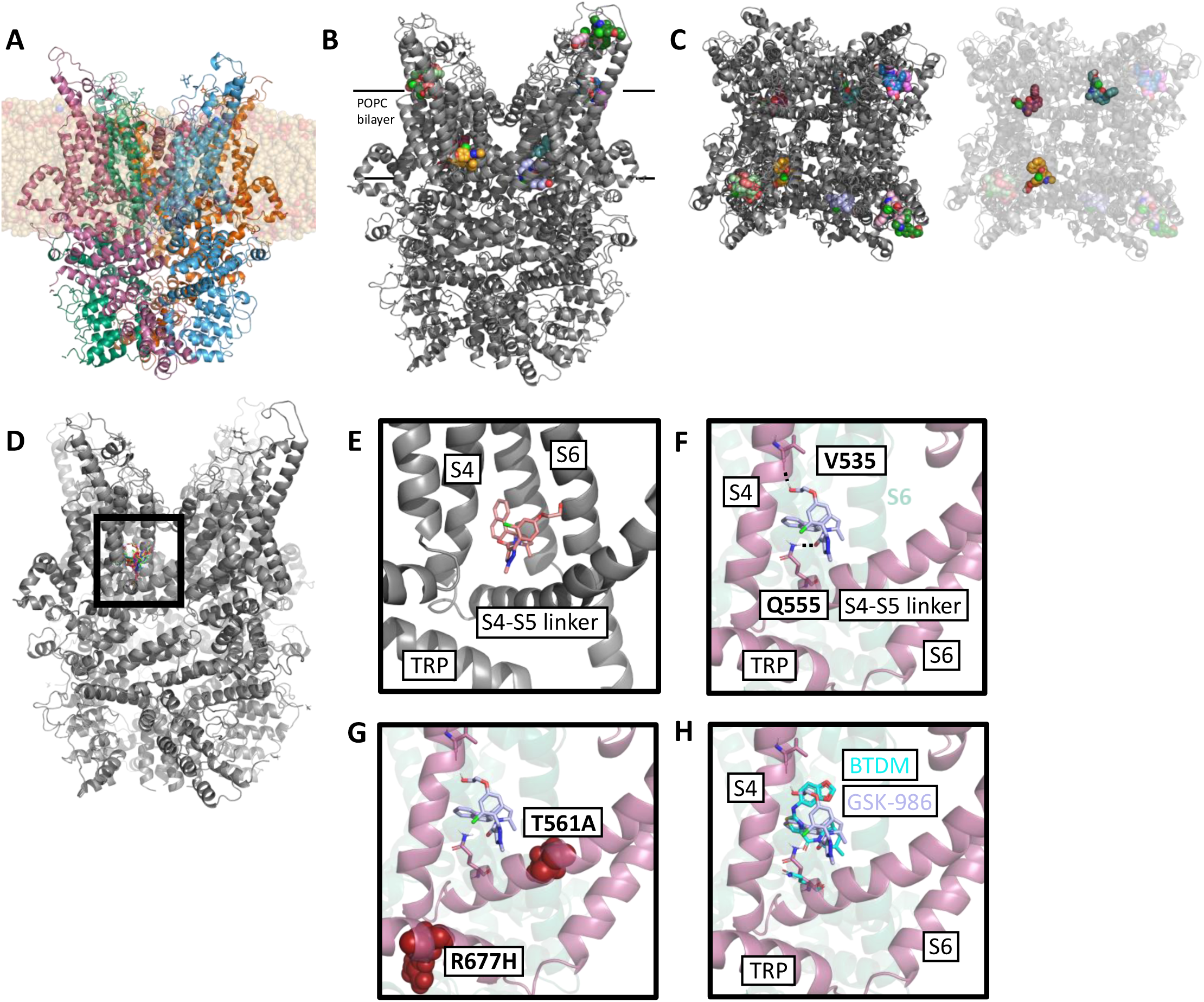
Docking and MD simulations predict the TRPC3 S4-S5 pocket as a potential site of binding for GSK-986. **(A)** TRPC3 structure PDB 6CUD embedded in a bilayer of the model lipid 1-palmitoyl-2-oleoyl-glycero-3-phosphocholin (POPC). **(B)** Top 20 results from Autodock Vina docking of GSK-986 against the whole protein from the final frame of 100-ns TRPC3 simulation (6CUD), with ligands shown as spheres. **(C)** Top-down (extracellular) view of top 20 docking poses. Three out of four S4/5 pockets are occupied by GSK-986. **(D)** Top 20 results from focused docking of GSK-986 against the S4-S5 pocket. **(E)** Pose chosen for further simulation. **(F)** The stable GSK-986 pose in the S4-S5 pocket, following equilibration, is in close contact to the sidechain of residue Q555, and the backbone of V535. **(G)** The pathogenic GOF mutations SCA41 (R677H) and *Mwk* (T561A) are also located in the S4-S5 region. **(H)** The predicted GSK-986 binding pose is analogous to the structurally resolved pose of the non-selective TRPC inhibitor BTDM in TRPC6 (cyan, PDB 7DXF). The equivalent glutamine residue in the S4-S5 linker in this structure is highlighted (cyan).

Due to agreement in top-scoring poses, high occupancy across the four subunits, and the known ligand-binding capacity of this pocket throughout the TRP superfamily (2), the S4-S5 pocket was assessed to be the most likely binding site for GSK-986 (Fig. 3D). We therefore proceeded with more directed docking focused on this binding pocket. Poses found during this process were in good agreement both with the initial poses and within themselves, and repeatedly centered around contact between the central carbonyl oxygen of the ligand and the S4-S5 linker residue Q555 (Figure S1D-F). The S4-S5 pocket is well-established as a site of ligand interaction in TRP channels; it is the site of capsaicin binding in TRPV1, and a frequent site of lipid interaction across the TRP superfamily (37, 38). Interestingly, the known TRPC3 GOF variants, i.e., the *Mwk* mouse mutation (p.T561A) and the SCA41 patient variant (p.R677H), also cluster around this site, providing further evidence of its importance in channel regulation (16, 17)(Fig. 3G). Moreover, the TRPC inhibitor BTDM is also known to bind to the S4-S5 pocket, forming an equivalent ligand carbonyl to Q555 interaction as that predicted for GSK-986 (Fig. 3H) (11). Together, these findings highlight the S4-S5 pocket as a site of TRPC3 regulation and support the physiological relevance of the docked GSK-986 poses.

However, given that docking scores have limited reliability, we next used MD simulation to evaluate the conformational stability of predicted binding poses. We observed a slight conformational shift in the equilibration phase (see methods) whereby the ‘tail’ of the compound moved from an interaction with the C_α_ of I641 on the S6 helix to an interaction with the V535 C_α_ on the S4 helix, which was mediated by rotation around the bond between carbon 12 and 13 of the compound (Fig. 2A). This orientation of the tail was then sustained across all NPT equilibrations and production runs (Fig. S1G). As such, we consider this slightly re-arranged pose as the most likely mode of interaction for GSK-986 with TRPC3. This pose was stable across three 100-ns simulations, throughout which the observed close contacts between GSK-986 and residues V535 and Q555 were significantly sustained (Fig. S1H, I).

### Functional validation supports the TRPC3 S4-S5 pocket as the site of GSK-986 interaction

To validate the computational predictions about the GSK-986 binding site, we employed site-directed mutagenesis to introduce a p.Q555A mutation into wildtype (WT) and SCA41 (R677H) TRPC3 constructs. This glutamine-to-alanine substitution disrupts the predicted key interaction between TRPC3 and the central carbonyl group of GSK-986. Whole-cell patch-clamp experiments indicated comparable current density between HEK293T cells transiently transfected with WT and Q555A TRPC3 plasmids (Fig. 4A). No significant difference in activation of both WT and mutant TRPC3 by GSK1702934A was observed (Fig. 4A-C). These results suggest that the Q555A mutation does not affect expression and/or pharmacologic activation of TRPC3.

**Figure 4.**
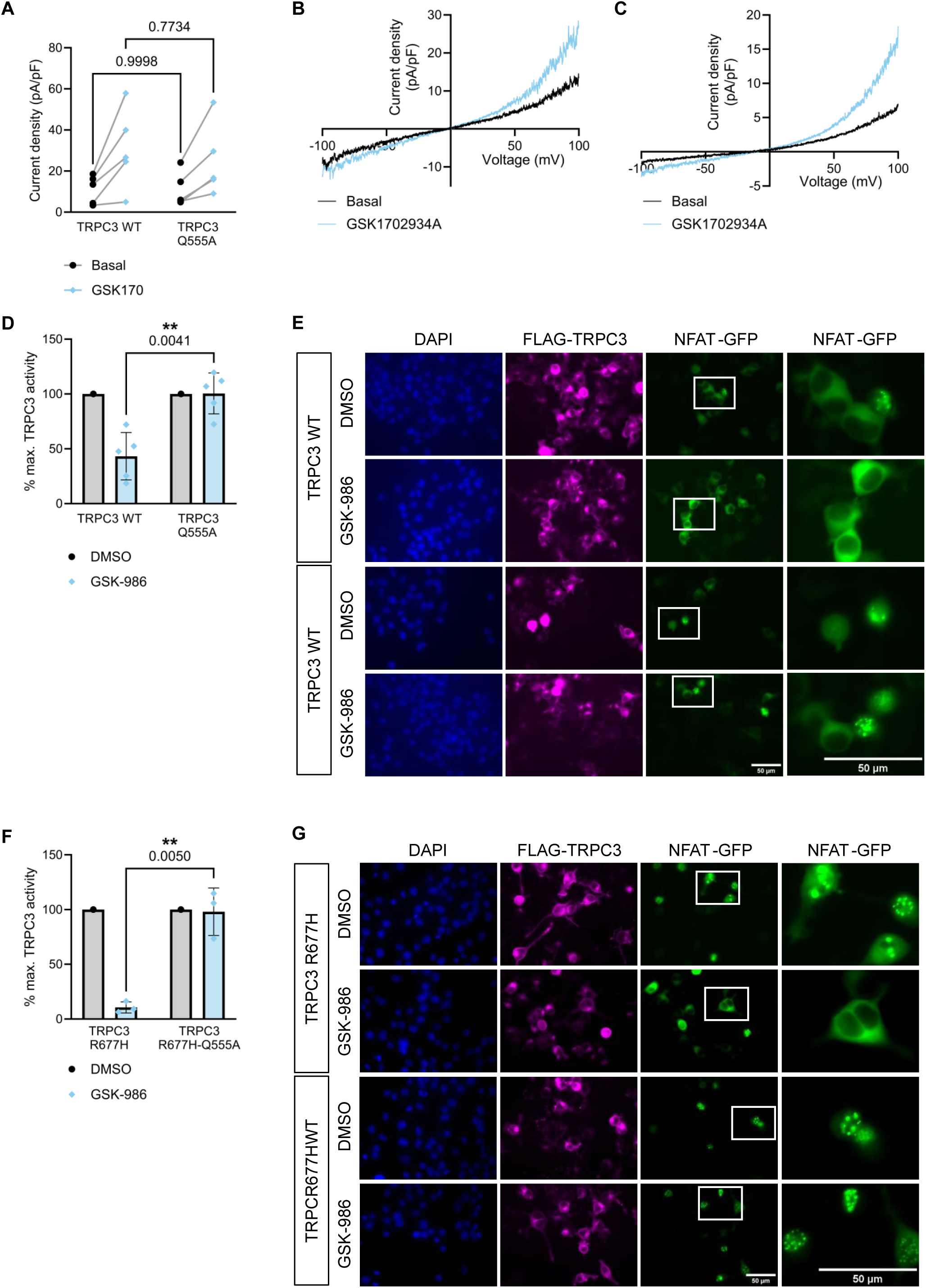
TRPC3 mutation Q555A impairs inhibition by GSK-986. **(A)** No significant difference in maximum and outward current density values was recorded using whole-cell patch-clamp between HEK293T cells transfected with TRPC3 wildtype (WT) and Q555A mutation constructs under both basal conditions and activation with 10 μM GSK1702934A (GSK170). Two-way ANOVA with Šídák’s multiple correction, **p=0.0041, n=5 cells. Scatter points represent biological replicates. Current density measurements were taken at 100mV, adjusted for baseline at a 0 mV hold. **(B, C)** Representative TRPC3 currents recorded from transiently transfected HEK293T cells using whole-cell patch-clamp. Cells were transfected with either WT TRPC3 (B) or the TRPC3 Q555A mutation construct (C). Basal traces were taken before activation with 10 μM GSK1702934A to compare with traces at peak activation, prior to desensitization. **(D)** Mutation Q555A significantly impairs the inhibition of TRPC3 by GSK-986 as demonstrated by impaired nuclear NFAT-GFP translocation in Neuro-2a cells. Cells were transiently transfected with WT or Q555A mutant FLAG-TRPC3 and treated with 10 μM of the activator GSK1702934A and either 1 μM GSK- 986 or an equivalent volume of DMSO. The percentage of nuclear GFP-NFAT of DMSO-treated cells was used as the maximum activation of TRPC3 and compared to inhibitor-treated cells. Two-way ANOVA followed by Šídák’s multiple comparisons test, **p=0.0041, n=5 independent biological replicates, each being the percentage of cells with nuclear-located GFP-NFAT across 50-200 cells. Scatter points represent biological replicates and bar graph shows mean ± standard deviation (SD). **(E)** Representative images of NFAT-GFP translocation experiments quantified in (D). Treated cells were fixed and immunostained with antibodies against FLAG and GFP. Nuclei are visualized with DAPI in blue. Scale bars: 50µm. Image windows are expanded to show NFAT localization in individual cells. **(F)** The Q555A mutation significantly impairs the GSK-986-mediated inhibition of TRPC3 harboring the GOF disease mutation R677H, as demonstrated by impaired nuclear NFAT-GFP translocation in Neuro-2a cells. Cells were transfected with FLAG-TRPC3 R677H with and without the additional Q555A mutation and treated with either 1 μM GSK-986 or an equivalent volume of DMSO. The percentage of nuclear GFP-NFAT of DMSO-treated cells was used as the maximum activation of TRPC3 and compared to inhibitor-treated cells. Two-way ANOVA followed by Šídák’s multiple comparisons test, **p=0.005, n=3 independent biological replicates, each being the percentage of cells with nuclear-located GFP-NFAT across 50-200 cells. Scatter points represent biological replicates and bar graph shows mean ± SD. **(G)** Representative images of NFAT-GFP translocation experiments quantified in (H). Treated cells were fixed and immunostained with antibodies against FLAG and GFP. Nuclei are visualized with DAPI in blue. Scale bars: 50µm. Image windows are expanded to show NFAT localization in individual cells.

We next used the NFAT-GFP translocation assay to determine the effect of the Q555A mutation on TRPC3 inhibition by GSK-986. We found that GSK-986 inhibited TRPC3 containing the Q555A mutation significantly less than WT TRPC3 (Fig. 4D, E). We also introduced the Q555A mutation into mutant TRPC3 harboring disease-causing GOF mutations. As with WT TRPC3 activated by GSK1702934A, the Q555A mutation significantly reduced GSK-986-mediated inhibition of both the SCA41 p.R677H variant (Fig. 4F, G) and the TRPC3 *Mwk* mutation (p.T561A) (Fig. S2). Together, the robust impairment of GSK-986 inhibition of TRPC3 by the Q555A mutation suggests that the S4-S5 pocket is indeed the site of binding for GSK-986.

In addition to the key residue Q555, five further residues in the S4-S5 pocket were identified as potential sites of contact with GSK-986 (Fig. 5A, B). To determine the role of these residues in inhibition of TRPC3 by GSK-986, we generated alanine mutations of each amino acid in the GOF TRPC3 R677H construct (W457A, S539A, L558A, G559A, and M574A) and used the generated constructs in the NFAT-GFP translocation assay. We found that only the L558A mutation significantly impaired GSK-986 inhibitor activity (Fig. 5C). As leucine has no active groups capable of polar interactions with GSK-986, it is possible that L558 forms hydrophobic contacts with the compound which stabilize the binding pose.

**Figure 5.**
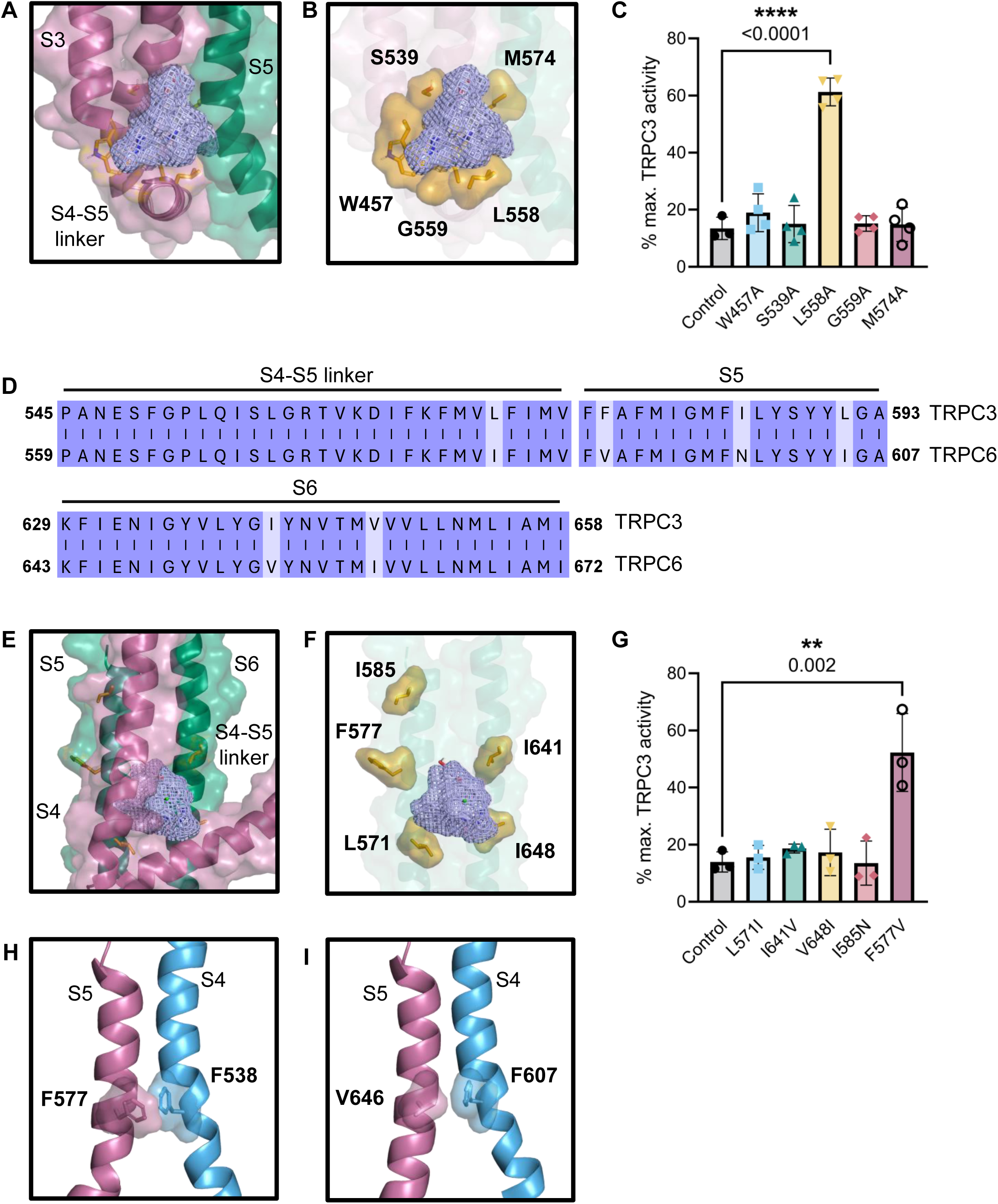
Residues L558 and F577 impact inhibition of TRPC3 by GSK-986. **(A)** GSK-986 (lilac mesh) in its stable interaction pose in the TRPC3 S4-S5 binding pocket, with close-contact residues highlighted in orange. **(B)** Five residues of interest form close contact with GSK-986. **(C)** NFAT-GFP translocation assay in Neuro-2a cells transfected with FLAG-TRPC3 harboring the GOF disease mutation R677H and additional alanine substitutions in residues of interest. Cells were treated with 1 μM GSK-986 inhibitor. Introduction of the L558A mutation significantly impairs the inhibition of NFAT-GFP nuclear translocation in GSK-986-treated cells. The percentage of nuclear GFP-NFAT of DMSO-treated cells was used as the maximum activation for each TRPC3 construct and to which inhibitor-treated cells were compared. Ordinary one-way ANOVA followed by Holm-Šídák’s multiple comparisons test, ****p<0.0001, n=3 independent biological replicates, each being the percentage of cells with nuclear-located GFP-NFAT across 50-200 cells. Scatter points represent biological replicates and bar graph shows mean ± standard deviation (SD). **(D)** High amino acid sequence conservation between TRPC3 and TRPC6 across the helices enclosing the S4-S5 pocket. **(E)** GSK-986 (lilac mesh) within the S4-S5 pocket with TRPC3 residues that differ in TRPC6 highlighted in orange. **(F)** Residues L571, F577, I585, I641, and I648 differ between the TRPC3 and TRPC6 sequence and are in close proximity to the GSK-986 binding site in the S4-S5 pocket. **(G)** NFAT-GFP translocation assay in Neuro-2a cells transfected with FLAG-TRPC3 harboring the GOF disease mutation R677H and additional TRPC3-to-TRPC6 substitutions in residues of interest. Cells were treated with 1 μM GSK-986 inhibitor. Introduction of the F577V mutation significantly impairs the inhibition of NFAT-GFP nuclear translocation in GSK-986-treated cells. The percentage of nuclear GFP-NFAT of DMSO-treated cells was used as the maximum activation for each TRPC3 construct and to which inhibitor-treated cells were compared. Lognormal ordinary one-way ANOVA followed by Holm-Šídák’s multiple comparisons test, **p=0.002, n=3 independent biological replicates, each being the percentage of cells with nuclear-located GFP-NFAT across 50-200 cells. Scatter points represent biological replicates and bar graph shows mean ± SD. **(H, I)** The TRPC3 F577 residue may have different functional effects on the channel compared to the equivalent valine residue in TRPC6. TRPC3 F557 appears to engage in perpendicular π-stacking with F538 on the adjacent S4 helix (H), while the TRPC6 V646 cannot (I).

Remarkably, despite the high sequence and structural homology between TRPC3 and TRPC6, GSK-986 displays a ten-fold selectivity for TRPC3 (Fig. 2B). Residues within and around the S4-S5 pocket are highly homologous between the two channels, consistent with an important functional mechanism for this region. There are only five residues around the predicted binding site that are different between TRPC3 and TRPC6 (Fig. 5D-F). We reasoned that one of these residues might contribute to the difference in GSK-986 activity against the two highly related channels. To investigate this, these five residues were mutated experimentally in the TRPC3 R677H mutant channel to their TRPC6 equivalents and then assessed for their impact on inhibition by GSK-986. Of these substitutions, F577V was found to significantly impair TRPC3 inhibition by GSK-986 (Fig. 5G). F577 appears to participate in π-stacking with F538 on the S4 helix in TRPC3, while this interaction is not present in TRPC6 at the equivalent V646 residue (Fig. 5H, I). This altered inter-subunit interaction may produce differences in channel dynamics that contribute to the observed selectivity for TRPC3 over TRPC6. The change in interaction between the S4 and S5 helix may also alter the size and shape of the S4-S5 pocket, affecting GSK-986 binding.

### TRPC3 conformational changes give insight into a potential mechanism of inhibition

Previous comparison of activator-bound and inhibitor-bound TRPC6 structures has shown a number of conformational differences, most notably in the S4-S5 linker helix (38). This comparison was performed by aligning the cryo-EM structure of TRPC6 bound to the activator AM-0883 (PDB 6UZ8), resolved near the DAG binding site, with cryo-EM structures of TRPC6 bound to the inhibitor AM-1473 (PDB 6UZA), resolved in the S1-S3 site, or inhibitor BTDM (PDB 5YX9), which is resolved in the vanilloid-like S4-S5 pocket. The authors observed that in both inhibitor-bound structures, the S4-S5 linker helix was maintained in an upwards configuration with respect to the plasma membrane. In contrast, the activator-bound TRPC6 S4-S5 linker helix displayed a lower, downward conformation. The authors attributed this conformational difference to a potential inhibitory mechanism, in which the S4-S5 linker is stabilized in the upwards configuration (38). In the case of AM-1473, stabilization of the S4-S5-linker was proposed to be mediated by an inhibitory lipid in the vanilloid site. This lipid has been resolved in cryo-EM structures of TRPC3 and TRPC6, including TRPC6-AM-1473 (PDB 6UZA), as well as other TRP channels (38). The reported conformational changes between activated and inhibited TRPC6 were not measured quantitatively but observed following alignment of the structures. To establish a quantitative measurement that represented the conformational change in the S4-S5 linker, we first reproduced this alignment using the same TRPC6 structures (Fig. S3A). Subsequently, based on the predicted interaction residues of GSK-986, measurements were taken between the C_α_ of the S4 valine (TRPC3:V535 and TRPC6:V604) and S4-S5 linker glutamine (TRPC3:Q555 and TRPC6:Q624) for multiple TRPC3 and TRPC6 structures (Fig. S3B). This served as an approximation of the S4-S55 pocket size and represented distance of the S4-S5 linker from the S4 due to an upward or downward conformation.

Alignment of the GSK-986-bound TRPC3 channel with the agonist-bound TRPC6 structure (TRPC6_AM-0883_) revealed that the inhibitor-bound TRPC3 subunit (TRPC3_986_) S4-S5 linker exhibited an upward conformation relative to the agonist-bound TRPC6 (Fig. 6A) consistent with the observations on inhibitor-bound TRPC6 (38). The ligand-free subunits (TRPC3_986(apo)_) showed much higher agreement with the slightly downward S4-S5 linker of the activator-bound TRPC6_6UZ8_ structure. Additionally, the alignment of TRPC3_GSK-986_ with the original cryo-EM structure used for the MD system setup (TRPC3_6CUD_) also showed a slight upward movement of the S4/5 linker, while TRPC3_apo_ showed higher agreement with the ligand-free, cryo-EM TRPC3_6CUD_ S4-S5 linker (Fig. 6B). Moreover, distance analysis between V353 C_α_ and Q555 C_α_ revealed that, across the simulations, the inhibitor-bound TRPC3_GSK-986_ subunit had a significantly smaller S4-S5 pocket compared to the TRPC3_apo_ subunits (Fig. 6C).

**Figure 6.**
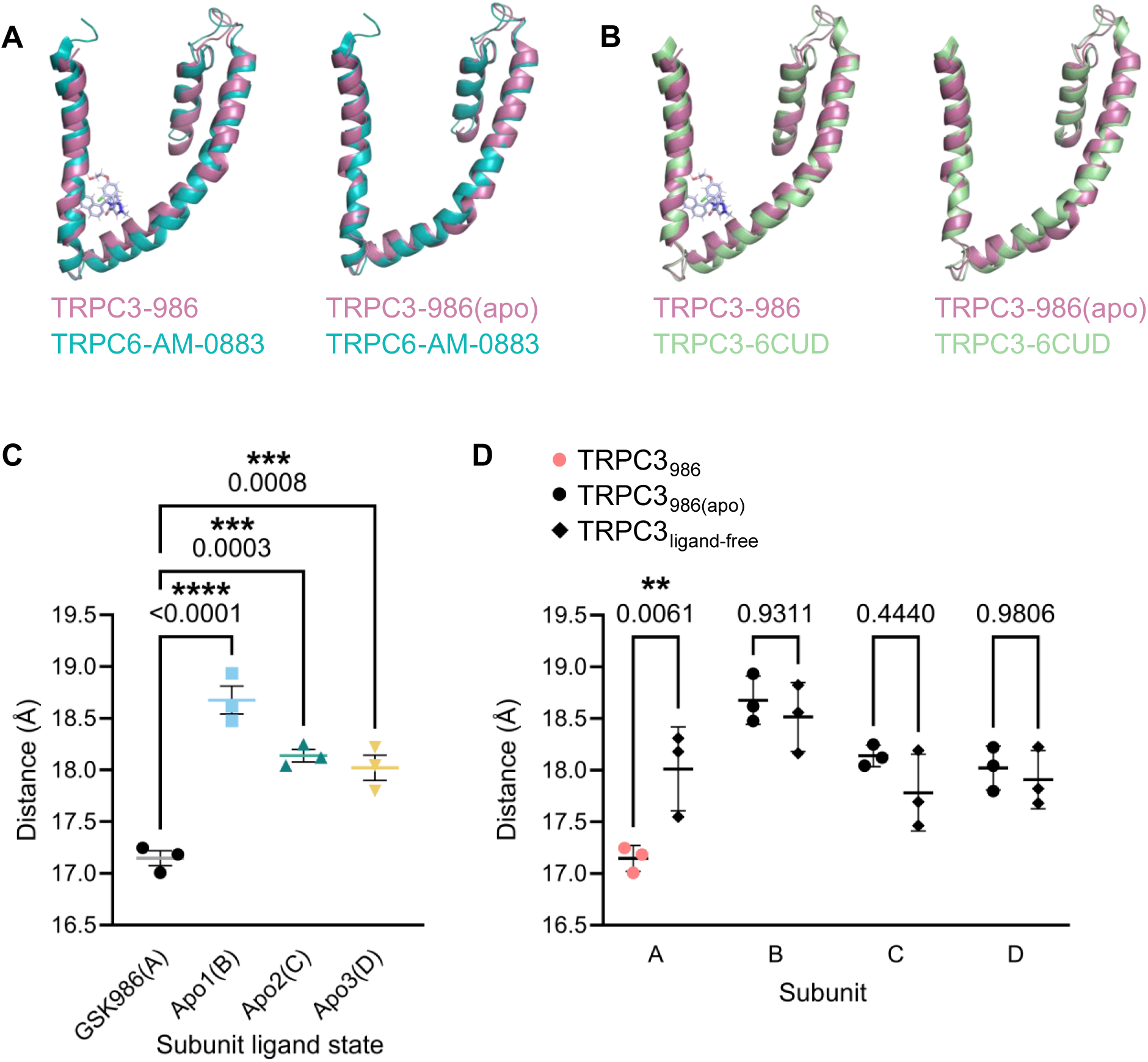
Binding of GSK-986 leads to a reduced S4-S5 pocket distance. **(A, B)** Alignments between the GSK-986-bound or apo subunit S4-S5 linkers of TRPC3 (light pink) with the activator-bound TRPC6 structure (TRPC6-AM-0883, 6UZ8, cyan) (A) or with the apo TRPC3 structure upon which simulations were based (TRPC3-6CUD, cyan) (B) show slight alterations in conformation. See also Figure S3. **(C)** The S4-S5 distance is significantly smaller in the GSK-986 simulation subunit occupied by the compound compared to any of the three apo subunits. Ordinary one-way ANOVA followed by Dunnett’s multiple comparisons test; ****p<0.0001, ***p=0.0003, ***p=0.0008, n=3 independent simulations. Scatter points represent independent replicates and bar graph shows mean ± standard deviation (SD). **(D)** The GSK-986 simulation subunit occupied by the compound (TRPC3_986_), subunit A, has a significantly shorter S4-S5 distance than its ligand-free simulation equivalent subunit A. In contrast, the apo subunit S4-S5 distances are not significantly shorter than their ligand-free counterpart subunits. Two-way ANOVA followed by Šídák’s multiple comparisons test, **p=0.0061, n=3 independent simulations. Scatter points represent independent replicates and bar graph shows mean ± SD.

To further demonstrate the difference in S4-S5 pocket size, measurements were compared individually between subunits A-D of the GSK-986-docked TRPC3 and ligand-free TRPC3 simulations. In GSK-986-docked TRPC3, the compound was only present in subunit A (TRPC3_986_), with subunits B, C, and D empty (TRPC3_986(apo)_) (Fig. 6D). The resulting measurements indicated that only the TRPC3_986_ subunit had a significantly shorter S4-S5 distance than its ligand-free counterpart throughout simulations, representing an S4-S5 linker in the upwards conformation. Together, these results support an inhibitory mechanism in which GSK-986 maintains the TRPC3 S4-S5 linker in an upward conformation relative to the S4 helix, consistent with the proposed mechanism for known inhibitors AM-1473, its analogue SAR7334, and BTDM (38–40).

### Impaired SAR7334 inhibition by TRPC3 Q555A supports proposed shared mechanism of inhibition

As with the compound BDTM, cryo-EM structures of the S4-S5 pocket lipid suggest an interaction with the same glutamine residue as we have identified for GSK-986 (Fig. 7A, B). As such, TRPC3 mutation Q555A may disrupt binding of the inhibitory lipid and impair inhibition by SAR7334 and AM-1473. To test this hypothesis, inhibition of SCA41 TRPC3 and SCA41-Q555A TRPC3 by SAR7334 was measured using the functional NFAT-GFP translocation assay in Neuro-2a cells. Inhibition of the SCA41-Q555A binding site mutant was significantly impaired compared to inhibition of SCA41 TRPC3 (Fig. 7C, D).

**Figure 7.**
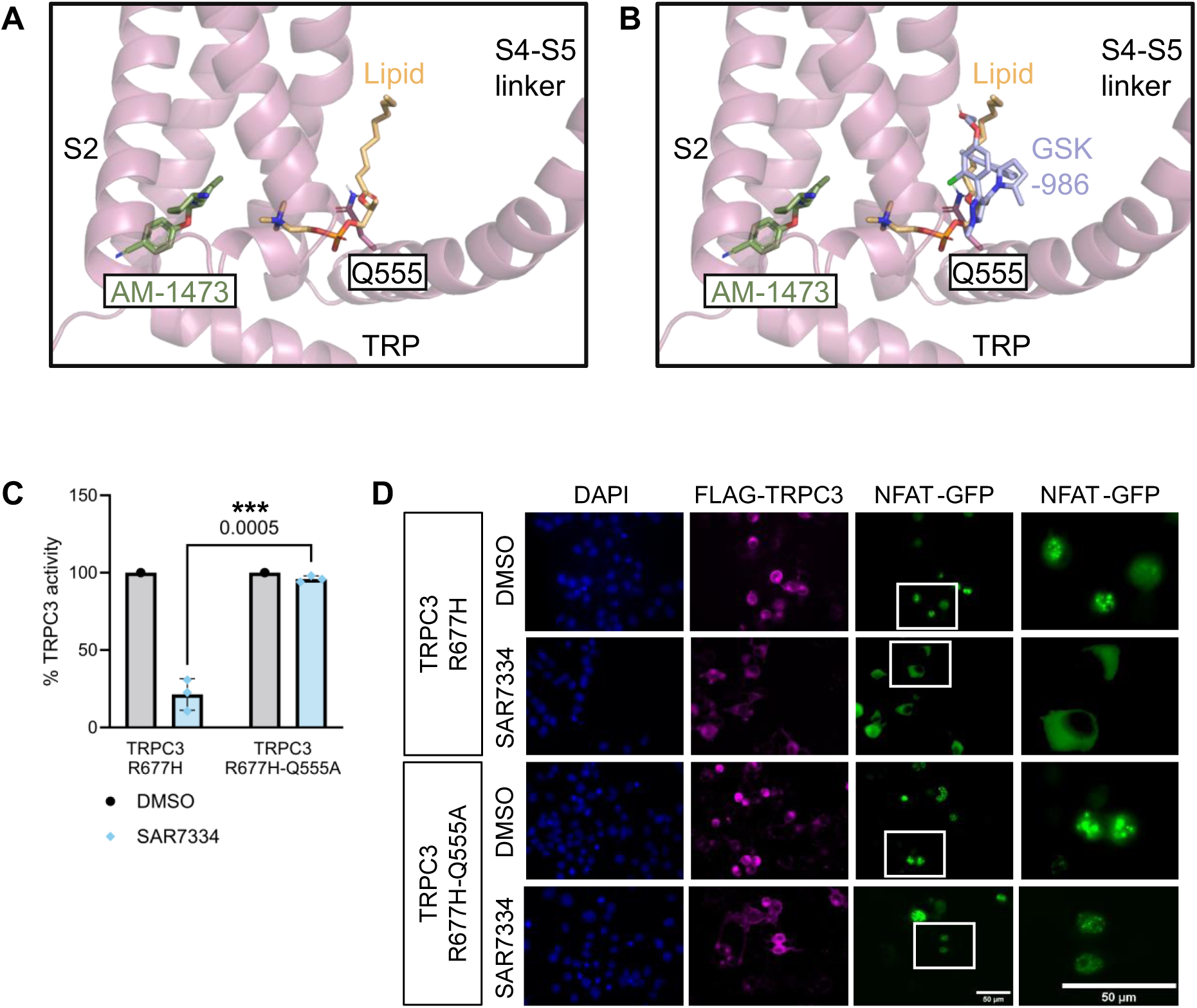
The S4-S5 pocket binding site mutation Q555A impairs inhibition by SAR7334. **(A)** TRPC6-AM-1473 (PDB 6UZ8) with SAR7334 analogue AM-1473 (green) in the S1-S3 pocket and the proposed inhibitory lipid (orange) in the S4-S5 pocket. The Q555 TRPC6 equivalent is shown as a potential interacting residue for the lipid. **(B)** TRPC6-AM-1473 with GSK-986 (purple) from TRPC3 simulations overlaid. Both GSK-986 and the lipid appear to be in contact with Q555. **(C)** Inhibition of SCA41 by SAR7334 is significantly impaired by mutation Q555A. Two-way ANOVA followed by Šídák’s multiple comparisons test, ***p=0.0005, n=3 independent biological replicates, each being the percentage of cells with nuclear-located GFP-NFAT across 50-200 cells. Scatter points represent biological replicates and bar graph shows mean ± standard deviation (SD). **(D)** Representative images of NFAT-GFP translocation experiments quantified in (C). Treated cells were fixed and immunostained with antibodies against FLAG and GFP. Nuclei are visualized with DAPI in blue. Image windows are expanded to show NFAT localization in individual cells. Background subtraction was performed equally across all images using ImageJ. Scale bars: 50 μm.

The combined evidence from molecular docking and simulation of GSK-986, mutagenesis of the binding site and surrounding residues, and the resolved structure of the similar, anilino-thiazole-derived TRPC3/6 inhibitor, BDTM, provide compelling support for the vanilloid-like S4-S5 pocket as the binding site for GSK-986. However, in light of the impaired inhibition by SAR7334 by TRPC3 mutation Q555A, further work was performed to exclude the S1-S3 pocket as an alternative site of GSK-986 binding. Docking of GSK-986 in the S1-S3 pocket was performed using QuickVina-W (36). The results showed little agreement in docked poses, and so the three top-scoring poses were selected to undergo 3 x 20-ns simulations to assess possible favorable binding positions. Four possible residues were highlighted as possible points of interaction for GSK-986 within this pocket (D461, H377, S439, and Y543). However, no significant impact on inhibition by the compound was observed in the NFAT translocation assay upon mutation to any of these residues to alanine (Fig. S4), suggesting that GSK-986 does not bind in this pocket. Together, these findings support the S4-S5 pocket as the binding site of GSK-986.

## Discussion

The development of specific TRPC3 inhibitors for experimental and clinical use has historically been challenging, in part, due to the high homogeneity of the TRPC3/6/7 subgroup (30). The relative lack of structural information on TRPC3 ligand binding and channel conformation, including the absence of any open-state TRPC structures, has also presented a challenge to understanding the mechanisms of TRPC3 activation and inhibition. Here, we have presented a novel, highly potent TRPC3 inhibitor, GSK-986, that exhibits ten-fold selectivity over close family member TRPC6. This compound is the most potent reported to date, with an IC_50_ of 0.08 nM, as measured by manual patch-clamp electrophysiology. Furthermore, using MD simulations combined with functional experiments, our work has identified the vanilloid-like S4-S5 linker as the likely site of binding for GSK-986 and has explored a potential structural mechanism by which this compound impairs activation of TRPC3.

The binding site we have identified for GSK-986 has previously been demonstrated as the binding site of a similar, anilino-thiazole-derived TRPC3/6 inhibitor, BTDM (40). Structural and functional data show that BTDM forms a comparable interaction to that predicted for GSK-986, between a central carbonyl group of the compound and the TRPC6 S4-S5 linker equivalent residue to the TRPC3 Q555. Our findings therefore support this pocket as a key site of TRPC3 and TRPC6 inhibition with implications for further development of specific antagonists.

In our simulations, GSK-986 binding stabilized the S4-S5 linker in an inhibitory conformation, in which the S4-S5 linker occupies a more upwards conformation in inhibitor-bound structures. This stabilization of the linker helix is consistent with our predicted binding pose. Throughout simulations, GSK-986 appears to form stable interactions with residues V535 and Q555 of the S4 and S4-S5 linker helices, which may couple the S4-S5 linker to the S4, resulting in the upward conformation (Fig. 3F). This is supported by structure-activity relationship data demonstrating that removal of the central carbonyl group eliminates all TRPC3 antagonism (GSK unpublished data). Based on general series SAR, removal of the entire ‘tail’ (see Fig. 2A) is expected to reduce TRPC3 antagonism. The central carbonyl group of the ligand is predicted to interact with the sidechain of Q555, and the OH-group off the ether group of GSK-986 with the backbone C_α_ of V535; loss of either of these interactions would prevent the proposed coupling of the S4 and S4-S5 linker. Further validation of this may be achieved by resolving TRPC3 structures under equivalent conditions in the closed, GSK-986-bound, and open-state conformation.

Interactions between residues of the S4-S5 linker and TRP helices have been implicated in the regulation of channel activation in several TRP channels, including TRPC3 (41–43). In TRPC3, disruption of a salt bridge between the S4/5-linker R560 (R572) and TRP helix E672 (E684), through either mutation R560E or E672R, reduces basal activity of the channel and impairs activation by GSK1702934A. Binding of PIP_2_ at an allosteric third lipid site (L3) was shown to regulate activation by GSK1702934A, as well as TRPC3 basal activity, with its effect mediated by the TRP and S4/5-linker helices (41). Building on the hypothesis described (38), we propose that stabilization of the S4-S5-linker by compounds such as GSK-986 may impair coupling of the TRP and S4-S5 linker helices and prevent propagation of activating conformational changes via the S4-S5 linker, resulting in inhibition of the channel (Fig. 8). This mechanism of inhibition may also explain the selectivity of GSK-986 for TRPC3 over TRPC6. The conformational changes between activator-bound and inhibitor-bound TRPC3 included movement of the S5 helix, as indicated in Figure 7. The altered inter-subunit interactions that may arise from the phenylalanine to valine substitution at position 577 of TRPC3 may impact the observed S5 conformational change and impair inhibition of TRPC6 by GSK-986, in addition to reducing inhibition of the TRPC3 F677V mutant channel (Fig. 5G).

**Figure 8.**
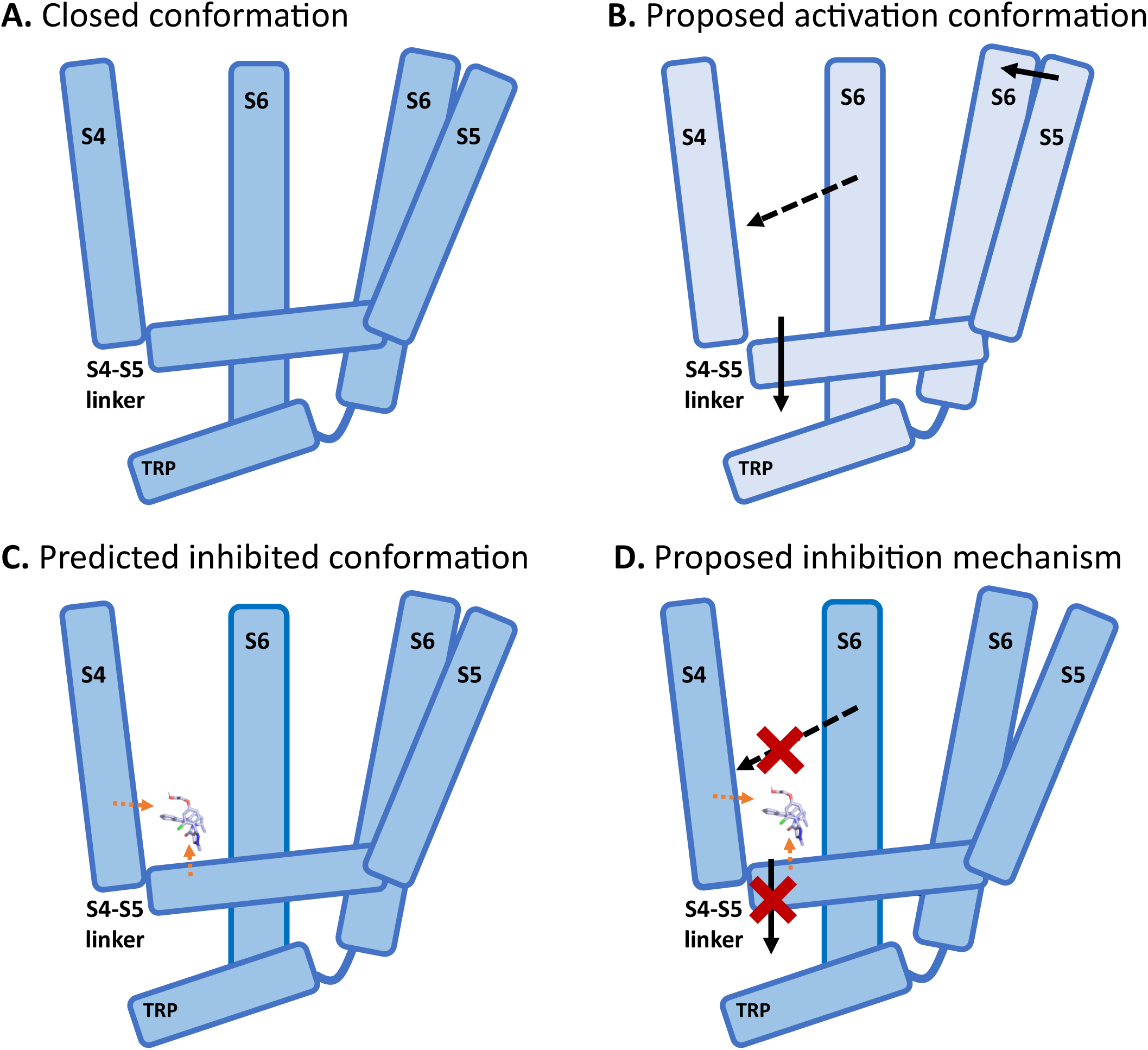
Proposed mechanism of inhibition by GSK-986 and other S4-S5 pocket TRPC3 inhibitors. **(A)** According to structural comparisons by Bai *et. al.* and molecular dynamics simulations presented in this study, the closed conformation (dark blue) of TRPC3 is proposed to have the S4-S5 linker in an upward conformation. **(B)** In the active conformation (light blue), the S4-S5 linker occupies a downward conformation. This is predicted to result in movement of the S5 and S6 helices, analogous to the role of the S4-S5 linker helix in coupling the voltage-sensing domain of the S4 linker to the pore-forming S5 and S6 in voltage-gated channels. The open conformation also likely requires movement of the S6 outward from the pore. **(C)** GSK-986 interacts with both the S4-S5 linker and the S4 helix. This likely couples the S4-S5 linker to the S4 and maintains the S4-S5 linker in the upward, inhibited conformation. Orange arrows indicate proposed protein-compound interactions. **(D)** The proposed mechanism of inhibition by GSK-986, and likely other S4-S5 pocket-binding inhibitors of TRPC3 and TRPC6, prevents the downward movement of the S4-S5 linker that is required for activation. Stabilization of the S4-S5 linker may effectively decouple the TRP and S4-S5 linker helices and may also impair outward movement of the S6 helix.

Overall, we have proposed the binding site for the novel, highly potent and selective TRPC3 inhibitor GSK-986 as the vanilloid-like S4-S5 pocket. Our work has reiterated the S4-S5 binding pocket as a key site of regulation in TRPC3, and supports a proposed mechanism of channel inhibition involving stabilization of the S4-S5 linker to prevent transition to the active-state conformation (38). These insights into inhibition of TRPC3 at the S4-S5 pocket are relevant for drug development and will inform future therapeutic strategies to combat the many pathologies linked to increased TRPC3 activity.

## Experimental procedures

### High-throughput Screening

#### Chemicals

All chemicals were obtained from Sigma unless otherwise stated. The tool compound GSK2820986A (GSK-986), a TRPC3/6 blocker, was identified by GSK in a previous high-throughput screening and optimization campaign and was provided by GSK for further study (34). All other tool compounds were purchased from Tocris.

#### Cellular Reagents

Unless stated otherwise, recombinant cells and BacMam viruses were generated by GSK. For TRPC3 and TRPC6 screening, HEK293 cells stably expressing MSRII (ATCC) or CHO-K1 cells stably expressing GAM-1 and EA1 were transduced with BacMam virus. For off-target profiling using automated patch clamp, hERG and K_V_1.5 channels were stably expressed in CHO-K1 cells, and Na_V_1.5 channels were stably expressed in HEK293 cells. For off-target profiling against Ca_V_1.2 using FLIPR, HEK293 cells stably expressing Ca_V_α2δ1 and β2a (GSK-Cytomyx Collaboration) were transduced with human α1C Ca_V_1.2 BacMam and IK1 BacMam.

#### FLIPR Assays

The FLIPR system (Molecular Devices) was used to assess drug effects by detecting changes in ionic activity within cells. TRPC3 and TRPC6 FLIPR data were previously generated by GSK (34). For Ca_V_1.2 studies, cells were transduced with BacMam, seeded at 15,000 cells/well into black, poly-d-lysine, clear-bottom 384-well plates, and incubated overnight at 37°C, 5% CO_2_. The following day, media was removed with a plate washer and replaced with FLIPR assay buffer (Tyrode’s Buffer supplemented with 20 mM HEPES, 11.9 mM NaHCO3, 2.5 mM probenecid, and 0.01% FAC Pluronic acid, pH 7.4). Cells were loaded with calcium-sensitive dye (FLIPR assay buffer supplemented with 2 µM Fluo4 and 250 µM Brilliant Black). Test compounds in FLIPR assay buffer with 30 mM final assay concentration (FAC) KCl were incubated for 15 min at 37°C. Inhibition of Ca_V_1.2 was assessed using depolarizing buffer (FLIPR assay buffer supplemented with 400 mM FAC KCl). Maximum RFU data were exported to ActivityBase^TM^ and normalized to controls for 4-parameter curve analysis.

#### Automated Electrophysiology

Cells were maintained in continuous culture for patch clamp recordings. Prior to experimentation, cells were seeded at 25,000 to 70,000 cells/mL in T175 cm² culture vessels and grown overnight at 37°C, 5% CO_2_ in air, unless stated otherwise. For TRPC3 studies, CHO-K1 cells stably expressing GAM-1 and E1A were transduced with BacMam (MOI ∼50) and incubated overnight with 1 µM TRPC3 inhibitor. For hERG studies, cells were incubated for 4 h at 37°C, then transferred to a 30°C incubator for 72 h. Patch clamp recordings were conducted using a 384-well IonWorks Quattro system (Molecular Devices) at room temperature in population patch clamp (PPC) mode. Cells were resuspended in DPBS containing Ca^2+^ and Mg^2+^, unless stated otherwise, at 2-3 million cells/mL. Calibration adhered to manufacturer guidelines for optimal seal resistance and capacitance compensation. Cells were seeded into the patch plate and allowed to seal for 4 min before perforation buffer (100µM FAC amphotericin in internal solution, 100 mM K-d-Gluconate, 50 mM KCl, 3.2 mM MgCl_2_, 10 mM HEPES, pH 7.3 with KOH, unless stated otherwise), was perfused and left to equilibrate for a further 9 min. Seal formation was monitored with a +10 mV pulse (100-200 ms) from the holding potential. Currents were acquired at between 1 and 10 kHz depending on the channel. Compound testing was performed in the presence 0.01% v/v Pluronic acid F-127. Wells with current responses <0.3 nA and seal resistances <30 MΩ were excluded. Data were exported to ActivityBase^TM^ and normalized to controls for 4-parameter curve analysis. For TRPC3 studies, a calcium-free external buffer was used (DPBS without divalents, supplemented with 1 mM MgCl_2_). A ramp voltage protocol was used to assess channel block in the presence and absence of compounds: holding potential of +10 mV, stepping to −100 mV for 50 ms, depolarizing to +100 mV over 200 ms, holding at +100 mV for 100 ms before returning to the holding potential. Test compounds were incubated for 1 min prior to agonist addition (10 µM GSK1702934A). Quasi-steady-state outward currents at +100 mV in the presence of agonist and test compounds were exported for analysis. For hERG studies, a chloride-based internal buffer was used (140 mM KCl, 1 mM MgCl_2_, 1 mM CaCl_2_, 20 mM HEPES, pH 7.3 with KOH). The following voltage protocol was used to assess channel block in the presence and absence of compounds: holding potential of −80 mV, stepping to −50 mV for 200 ms, +40 mV for 5000 ms, −50 mV for 5000 ms before returning to −80 mV. Test compounds were incubated for 4 min. hERG tail current were measured at −50 mV inactivation step (maximum response). For Na_V_1.5 studies, cells were clamped at −80 mV, measuring ionic currents with the protocol: test potential of 0 mV for 200 ms then back to holding potential, applied 5 times at 2 Hz. Test compounds were incubated for 10 min. Na_V_1.5 peak currents were measured at 0 mV activation step (minimum response). For K_V_1.5 studies, cells were clamped at −70 mV, measuring ionic currents with the protocol: test potential of +40 mV for 200 ms, 0 mV for 50 ms, then back to holding potential, applied 10 times at 2 Hz. Test compounds were incubated for 10 min. Na_V_1.5 peak currents were measured at 0 mV activation step (minimum response).

### Plasmids

The 3xFLAG-TRPC3, and 3xFLAG-*Mwk*TRPC3 have previously been described (44). Point mutations were generated using site-directed mutagenesis (QuickChange XL II, Agilent Technologies). Constructs were verified using Sanger sequencing (Source Biosciences). The CD8 plasmid used for transfections for electrophysiology was a kind gift from the Tammaro lab.

### NFAT-GFP Translocation Assay

Neuro-2a cells (American Type Culture collection, ATCC) were grown under standard conditions, in high-glucose Dulbecco’s Modified Eagle Media (DMEM) supplemented with 10% fetal bovine serum (FBS) and 1% Penicillin-Streptomycin. Cells were maintained at 37°C and 5% CO_2_.

Transfections were performed in Opti-Mem (Gibco) using FuGENE HD Transfection Reagent (Promega), according to manufacturer’s instructions, using a total of 0.5 µg DNA per well and a 3:1 ratio of FuGENE to DNA. Cells were co-transfected with 3xFLAG-TRPC3 (0.4 µg) and HA-NFAT-GFP (0.1 µg) 24 h after plating. Cells were fixed 24 h post-transfection with 4% paraformaldehyde (PFA) for 20 minutes, followed by membrane permeabilization in TBST with 0.1% Triton-X. Cells were blocked in a 10% skimmed milk and 1% normal goat serum (NGS) TBST solution, before immunostaining with antibodies for GFP (1:1,000, Takara Bio Clontech, Cat#632592) and FLAG (1:1,000, Sigma, Cat#F3165), followed by fluorescent secondary antibodies (Alexa-488, α-rabbit (Cat#A-11034); Alexa-594, α-mouse (Cat#A-11032); both 1:1,000, Invitrogen). Nuclei were stained using a DAPI-containing VectaShield medium (Vector Laboratories). Cytoplasmic vs nuclear NFAT-GFP localization was measured in 50-200 cells per condition, unless otherwise stated. Cells were obtained from a minimum of three independent experiments, using cells from different starting culture passages. Data were analyzed and graphs produced using Prism v9 (GraphPad). Images were processed using Fiji ImageJ (45).

### Whole-cell patch-clamp of HEK293T cells

HEK293T cells (American Type Culture collection, ATCC) were used for whole-cell patch-clamp electrophysiology. For dose-response experiments of GSK-986 inhibition of WT TRPC3 (Fig. 2D, E), cells were co-transfected in tissue-culture-treated plastic dishes with 3xFLAG-TRPC3 (2.5 µg) and CD8 (0.1 µg). Transfections were performed using FuGENE at a 3:1 ratio with DNA. Cells were then split and seeded onto glass coverslips at 18-24 h post-transfection and used for patch-clamp experiments. Independent replicates were taken as recordings from independent individual cells. For experiments assessing the activation of WT TRPC3 and TRPC3 Q555A by GSK1702934A (Fig. 4B, C), cells were seeded on glass coverslips followed by co-transfection with 3xFLAG-TRPC3 or 3xFLAG-TRPC3-Q555A (0.25 μg) and CD8 (0.01 μg). Transfections were performed using FuGENE at a 3:1 ratio with DNA, followed by media replacement at 4 h post transfection with fresh culturing media. Cells were used for patch-clamp experiments from 20 h post transfection. CD8-positive cells were identified under the microscope using superparamagnetic beads coated in monoclonal anti-CD8 antibody (Dynabeads^TM^ CD8, Thermo Fisher Scientific), which were added to cells before transferring to the bath.

Recordings were performed at room temperature using micropipettes pulled from thin-walled, filamented borosilicate glass capillaries and fire-polished, that gave resistances of 2- 3.5 MΩ in the experimental solutions. Currents were recorded using an Axon^TM^ Axopatch^TM^ 200B (Molecular Devices, UK) amplifier, with 2 kHz low-pass filtering. Currents were digitized at 10 kHz using Axon^TM^ Digidata® 1440A (Molecular Devices, UK). An Ag-AgCl reference electrode was used, connected to the bath solution via an agar salt bridge. Data was acquired using Axon^TM^ pCLAMP^TM^ v.11 (Molecular Devices, UK) and analyzed using Clampfit (Molecular Devices, UK) and Origin Pro (OriginLab®).

For all recordings, a voltage ramp was applied at 1 Hz at room temperature. Voltage was stepped from a holding potential of 0 mV to −100 mV for 100 ms, ramped to + 100 mV over 500 ms, held at +100 mV for 100 ms and stepped back to the holding potential. Series resistance was <10 MΩ and recordings were compensated 65-80%. For dose-response experiments, compounds were applied to the bath using gravity perfusion. TRPC3 was maximally activated by 1 µM GSK1702934A, and increasing concentrations of GSK-986 were co-applied. The mean current amplitude at +100 mV was measured and antagonist dose-response curves were obtained based on the following parameters: Y=Bottom + (Top-Bottom)/(1+10^((LogIC50-X)*HillSlope)), using Prism v9 (GraphPad).

Nominally Ca^2+^-free extracellular and intracellular solutions were used to minimize TRPC3 desensitization. An extracellular bath solution was used as follows (in mM): 140 NaCl, 5 KCl, 10 EGTA, 1 MgCl_2_, 10 glucose, 10 HEPES, pH 7.4 with NaOH; and the intracellular solution composition was (in mM): 135 CsCl, 5 EGTA, 5.5 MgCl_2,_ 10 HEPES, 5 Na_2_ATP, 0.1 Na_2_GTP, pH 7.4 with CsOH. For activation experiments of WT TRPC3 and TRPC3 Q555A, Ca^2+^-containing solutions were used to compare a more physiological response of the TRPC3 constructs. The extracellular solution was as follows (mM): 140 NaCl, 6 KCl, 2 CaCl_2_, 1 MgCl_2_, 10 glucose, 10 HEPES, pH 7.4 using NaOH; and the intracellular: 135 CsCl, 5 EGTA, 1 CaCl_2_, 1 MgCl_2_, 10 HEPES, 5 Na_2_ATP, 0.1 Na_2_GTP, pH 7.2 using CsOH.

### TRPC3-bilayer system preparation for molecular dynamics simulations

*6CUD TRPC3 system for GSK-986 MD simulation*: PDB ID 6CUD (10) was used as the starting point for docking and MD simulations. This structure is missing residues 1-21, 281-291, and 688-757. The loop region 281-291 was modelled using MODELLER (v.10.1) (46). Region 658-757 was not modelled. Instead, ends were capped with N-Methyl (NME) and acetyl (ACE) groups using PyMOL (Schrödinger). The missing N-terminal was also capped with an ACE group, and incomplete residues were modelled using SPDBV (47). This structure was parameterized with the AMBER99SB-ILDN forcefield (48). The channel was embedded in a homogenous bilayer of 615 1-palmitoyl-2-oleoyl-*sn*-glycero-3-phosphocholine (POPC) lipids, built in Martini (49) and converted to atomistic structure using *CG2AT.py* (50). The protein was embedded in this bilayer using a modified *InflateGro.py* script. The system was hydrated and neutralized with 150 mM NaCl. β-D-N-acetylglucosamine glycosylation at asparagine 404 was modelled using DoGlycans (51). This package allows processing of sugar residues by AMBER forcefields.

The system underwent <5000 steps of fastest descent energy minimization to reach a maximum force (F*_max_*) of 1000 kJ/mol/nm^2^. This was followed by equilibration under constant-temperature, constant-volume conditions (the canonical, NVT ensemble) at 310 K for 100 ps. Multiple rounds of constant-temperature, constant-pressure equilibration (NPT ensemble) were then performed prior to production runs, consisting of one 40-ns NPT equilibration with Cα position restraints increased step-wise from 100-1000 kJ/mol/nm^2^, followed by a 40-ns NPT equilibration with position restraints removed step-wise from 1000-100 kJ/mol/nm^2^, and then a final 10-ns NPT equilibration with position restraints gradually removed, from 100-0 kJ/mol/nm^2^. To generate independent starting points for repeat simulations, one further 10-ns NPT equilibration was performed per additional repeat with constant restraints of 100 kJ/mol/nm^2^. Further equilibration of the system was performed after addition of the docked GSK-986 poses, leading to removal of one POPC due to atomistic clashes. This resulted in a final bilayer of 614 POPC lipids for systems containing GSK-986 in either the S1-S3 or S4-S5 site.

### Ligand preparation

The 3D GSK-986 ligand structure was created using Marvin Sketch (ChemAxon, 2019). The 3D conformation was generated using Open Babel 2.4.1 (52). AutoDockTools was used to prepare the ligand file for docking by merging non-polar hydrogens (53). AM-BCC charges were generated using Antechamber (54) and topology files for use in the MD simulations generated using ACPYPE (55). The ligand was parameterized using GAFF (the Generalized Amber Force Field) (56).

### Docking

Undirected docking was performed using QuickVina-W (36). Once sites had been established, directed docking was performed to generate binding poses. For simulation, the GSK-986-containing system underwent minimization for <5,000 steps using a steepest descent algorithm, and 100-ps NVT ensemble equilibration. Independent 10-ns NPT ensemble equilibrations were performed for each of the three production runs, with constant restraints of 100 kJ/mol/nm^2^.

### Simulation details

All protein structures were parameterized using the AMBERff99SB-ILD forcefield (48) and default protonation states assigned during *pdb2gmx* processing in GROMACS (57). All systems used the TIP3P water model (58) and the lipid model used was SLipids (59). Systems were confined to rectangular boxes, solvated, and neutralized with 150 mM NaCl. During all equilibration and production, a timestep of 2 fs was used, and the pressure and temperature maintained at 1 bar and 310 K, respectively. During NVT ensemble equilibration, temperature was maintained through V-rescale coupling (60) with protein and non-protein coupling groups. During NPT ensemble equilibration, temperature was maintained through V-rescale coupling, with protein and membrane and water and ions coupling groups, and pressure was maintained through Berendsen pressure coupling (61). For production simulation, temperature was maintained in the same manner as the NPT equilibration, and pressure was maintained through the Parrinello-Rahman barostat (62). In all simulations, the LINCs algorithm (63) was used to constrain bonds between heavy atoms and hydrogen atoms. For short range LJ interactions, a cut-off of 1 nm was used, and particle mesh Ewald (PME) (64) was used for long range interactions. Periodic boundary conditions were imposed.

### Computational analysis

Trajectory data was analyzed using MDAnalysis 1.0 and 2.0 (65, 66) and Python. Trajectories were modified using GROMACS function *trjconv* and visualized using VMD (v.1.9.3) (67) and PyMOL (v.2.6) (68). All Python code used Python 3.6 (69)(Vand was written using PyCharm (v.2019.3.4) (JetBrains, 2017), using packages Numpy (70) and pandas (71). Graphs were plotted using Matplotlibs (72), Seaborn (73), and Prism v9 (GraphPad). OpenBabel (v.3.1.0) (74) was used to convert file formats when needed. Statistical tests were performed as outlined, and normality and lognormality were tested for using Shapiro-Wilk test; statistical analysis was performed using Prism v9 and Prism v10 (GraphPad).

## Data availability

Original data are available upon request from the corresponding authors. MD data (trajectories and input files) are uploaded to zenodo.org (DOI: 10.5281/zenodo.18151950).

## Supporting information

This article contains supporting information. Bai, Y., Yu, X., Chen, H., Horne, D., White, R., Wu, X., Lee, P., Gu, Y., Ghimire-Rijal, S., Lin, D.C.-H., Huang, X., 2020. Structural basis for pharmacological modulation of the TRPC6 channel. eLife 9, e53311. https://doi.org/10.7554/eLife.53311

## Author contributions

Conceptualization, E.B.E.B., P.C.B., J.C.; Data curation, E.B.E.B., P.C.B., J.C., B.A.C., K.M.G.; Formal analysis, J.C., B.A.C., Funding acquisition, E.B.E.B., P.C.B., J.C.; Investigation, J.C., B.A.C., Methodology, E.B.E.B., P.C.B., J.C., B.A.C., K.M.G.; Project administration, E.B.E.B., P.C.B., J.C.; Resources, E.B.E.B., P.C.B.; Supervision, E.B.E.B., P.C.B.; Visualization, E.B.E.B., J.C.; Writing – original draft, E.B.E.B., J.C.; Writing – review & editing, E.B.E.B., P.C.B., J.C., B.A.C., K.M.G., J.L.

## Acknowledgments

We thank the GSK Discovery Partnerships with Academia Collaboration (DPaC) project team and contributors including, Tony Dean (Chemistry lead, former Department of Chemical Sciences, GSK), Richard Kasprowicz (Discovery Biology & Screening, GSK), screening team (Discovery Biology & Screening, GSK) and subsequent support from David Washburn (Discovery Chemistry, GSK). We thank Stephen Tucker at the University of Oxford for providing electrophysiology equipment and Paolo Tammaro for providing the CD8 plasmid.

## Funding and additional information

This work was funded by Wellcome Trust Grant 219912/Z/19/Z, as part of 102161/B/13/Z.

