## supporting information for "Structural mechanism of TRPC3 inhibition by a potent and selective antagonist"

List of included material: Figures S1-S4

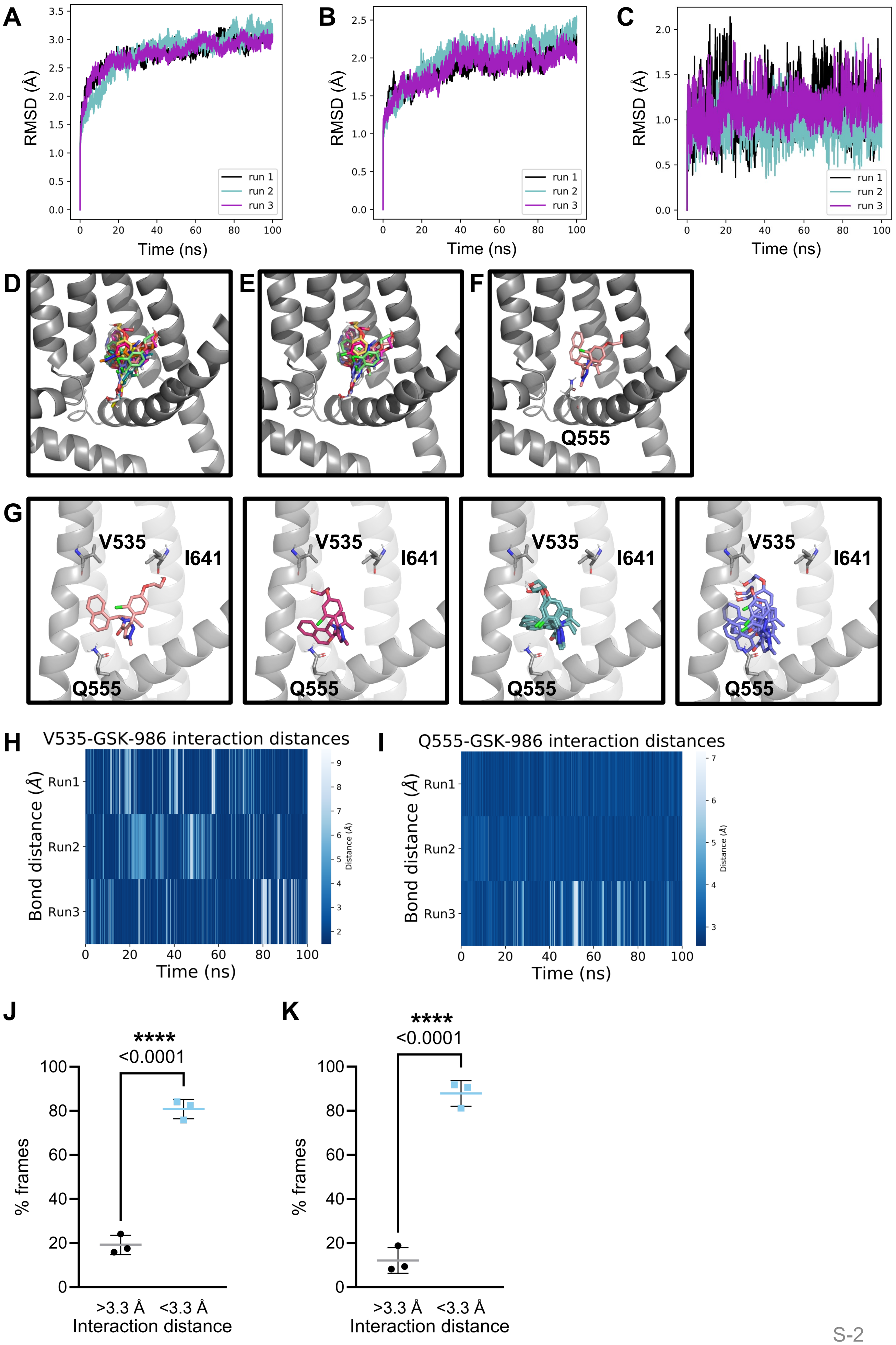
A
B
C
RMSD (Å)
Time (ns)
run 1
run 2
run 3
D
E
F
Q555
G
V535
I641
Q555
H
V535-GSK-986 interaction distances
Bond distance (Å)
Run1
Run2
Run3
Distance (Å)
Time (ns)
I
Q555-GSK-986 interaction distances
J
K
****
<0.0001
% frames
>3.3 Å
<3.3 Å
Interaction distance

**Figure S1. Docking and MD simulations predict the S4-S5 pocket as a potential site of binding for GSK-986. (A)** Root-mean-squared deviation of protein backbone atoms over 3 x 100-ns simulations of unbound TRPC3. **(B)** RMSD of protein backbone atoms over 3 x 100-ns simulations of GSK-986 docked in the S4-S5 pocket. **(C)** RMSD of GSK-986 atoms over 3 x 100-ns simulations. **(D)** Top ten GSK-986 poses docked to the S4-S5 pocket. **(E)** S4-S5-docked poses featuring contact between the ligand carbonyl oxygen and Q555. **(F)** Q555-contacting S4-S5 pose chosen for further simulation. **(G)** The initial pose is stable throughout energy minimisation (salmon), followed by immediate rearrangements of the ligand tail and double ring during NVT equilibration (pink). The output conformation is largely maintained across three independent NPT equilibrations, with minor rearrangements of the double ring (teal). The ligand conformation is largely stable across 3 x 100-ns unrestrained simulations (purple). **(H)** Distance analysis between the GSK-986 carbonyl oxygen and Q555 side chain nitrogen, in angstroms, over 3 x 100-ns simulations. **(I)** Distance analysis between the GSK-986 tail OH group and backbone oxygen of V535, in angstroms, over 3 x 100-ns simulations. **(J)** The distance between the GSK-986 carbonyl oxygen and Q555 side chain nitrogen is significantly below 3.3 Å across 3 x 100-ns simulations. Unpaired T-test, n = 3 independent simulation trajectories, ****$p<0.0001$, shown as mean ± SD. **(K)** The distance between the GSK-986 tail OH group and V535 backbone oxygen is significantly below 3.3 Å across 3 x 100-ns simulations. Unpaired T-test, n = 3 independent simulation trajectories, ****$p<0.0001$, shown as mean ± SD.

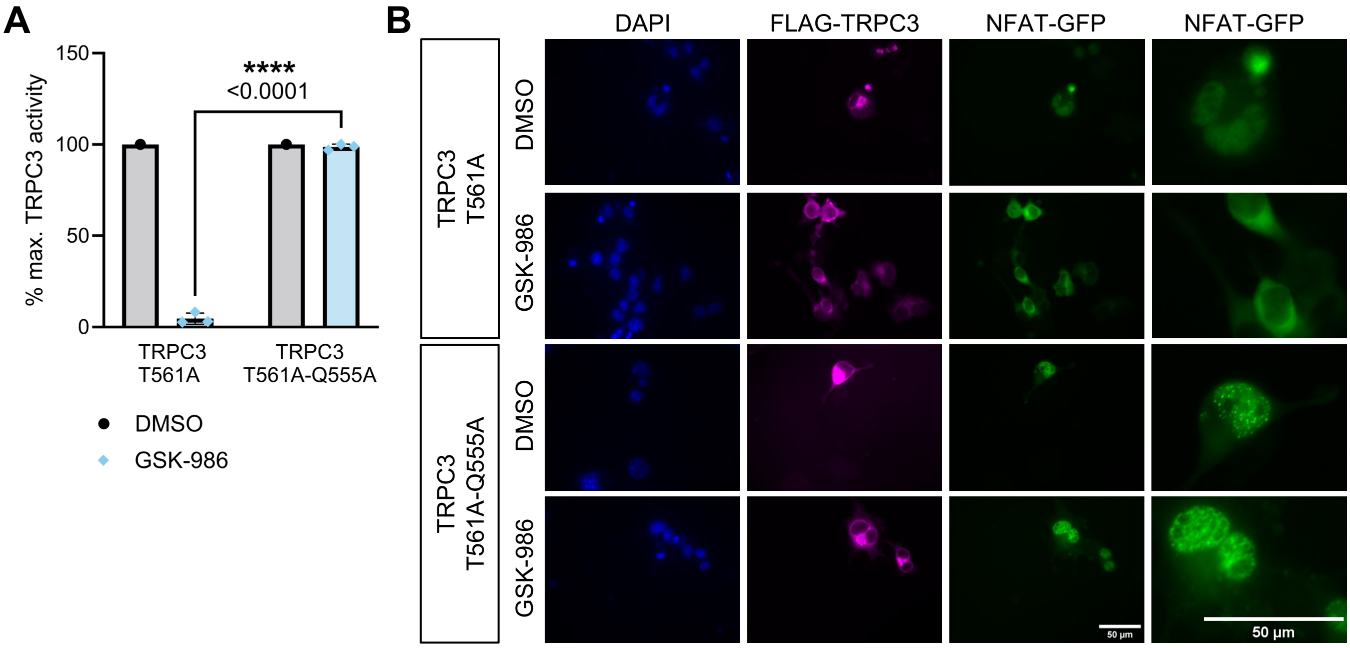


**Figure S2. Mutation Q555A impairs GSK-986 inhibition of TRPC3 harboring the gain-of-function *Mwk* mutation. (A)** The Q555A mutation significantly impairs the GSK-986-mediated inhibition of TRPC3 harboring the gain-of-function disease mutation T561A, as demonstrated by impaired nuclear NFAT-GFP translocation in Neuro-2a cells. Cells were transfected with FLAG-TRPC3 T561A with and without the additional Q555A mutation and treated with either 1 μM GSK-986 or an equivalent volume of DMSO. The percentage of nuclear GFP-NFAT of DMSO-treated cells was used as the maximum activation of TRPC3 and compared to inhibitor-treated cells. ****$p<0.0001$. Two-way ANOVA followed by Šídák's multiple comparisons test, n=3 independent biological replicates of ≥25 cells each. GSK-986 data shown as mean ± SD. **(B)** Representative images of NFAT-GFP translocation experiments quantified in (A). Treated cells were fixed and immunostained with antibodies against FLAG and GFP. Nuclei are visualized with DAPI in blue. Image windows are expanded to show NFAT localization in individual cells. Scale bars: 50μm.

**A**

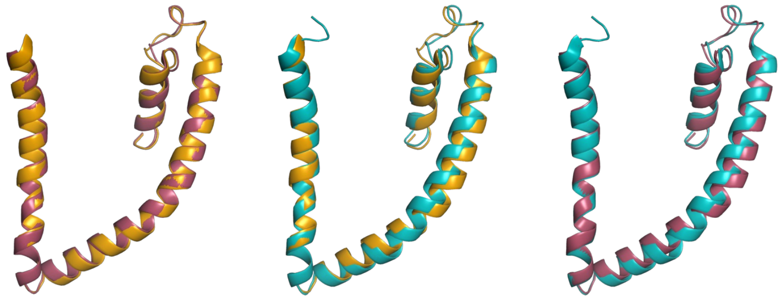


**B**

| Structure | Ligand | V535-Q555 distance (Å) |
|---|---|---|
| TRPC6$_{5XY9}$ | BTDM*, inhibitor | 18.4 |
| TRPC3$_{6UZA}$ | AM-1473, inhibitor | 18.6 |
| TRPC3$_{6UZ8}$ | AM-0883, activator | 19.2 |
| TRPC3$_{6CUD}$ | Apo** | 19.2 |
| TRPC3$_{986}$ | GSK-986, inhibitor | 17.15 (± 0.13) |
| TRPC3$_{986(apo)}$ | Apo | 18.28 (± 0.16) |
| TRPC3$_{ligand\text{-}free}$ | Apo | 18.05 (± 0.42) |

**Figure S3. Inhibitor-bound TRPC3 structures adopt an S4-S5 linker conformation that results in shorter S4-to-S4-S5 distances. (A)** Comparison of S4/5-linker conformation between inhibitor-bound TRPC6 structures (TRPC6-AM-1473 and BTDM, PDB 6UZA and 5YX9) with activator bound (TRPC6-AM-0883, PDB 6UZ8). Inhibitor-bound S4-S5 linker conformations occupy a slightly upwards conformation compared to the activator-bound, as reported (Bai et. al.). **(B)** Table reporting S4-S5 distances for TRPC6 and TRPC3 structures, as measured between the Cα atoms of V535 and Q555, or equivalent residues, as well as simulation subunits. *TRPC6 structure 5XY9 was resolved with compound BTDM, but the ligand was not resolved to atomic resolution. **TRPC3 structure 6CUD was resolved with a likely non-functional lipid in the diacylglycerol (DAG) binding site but is a closed-state structure and was not resolved in the presence of other ligands.

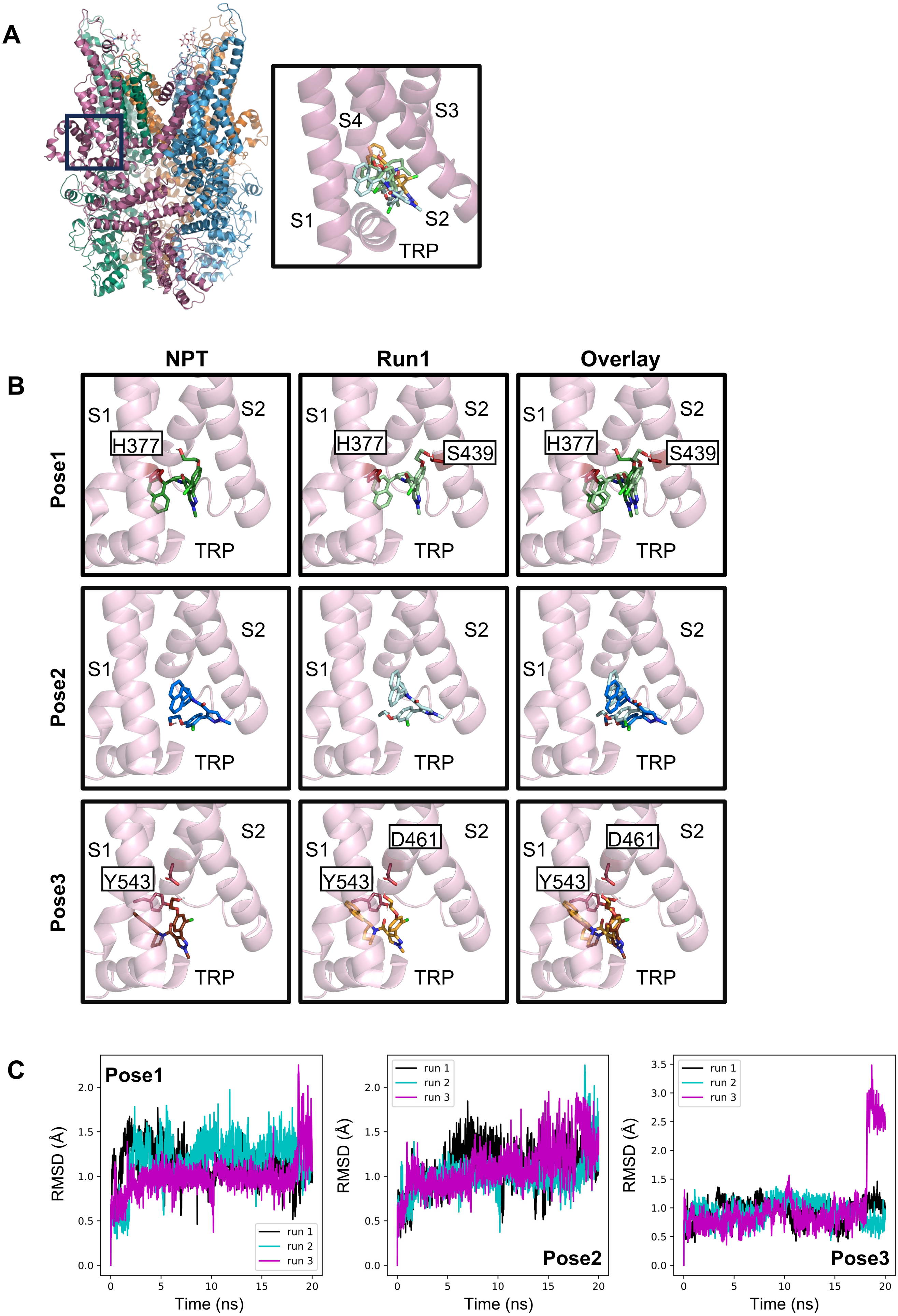

A
S4
S3
S1
S2
TRP
B
NPT
Run1
Overlay
Pose1
Pose2
Pose3
S1
S2
H377
S439
Y543
D461
TRP
C
Pose1
Pose2
Pose3
run 1
run 2
run 3
RMSD (Å)
Time (ns)

**D**

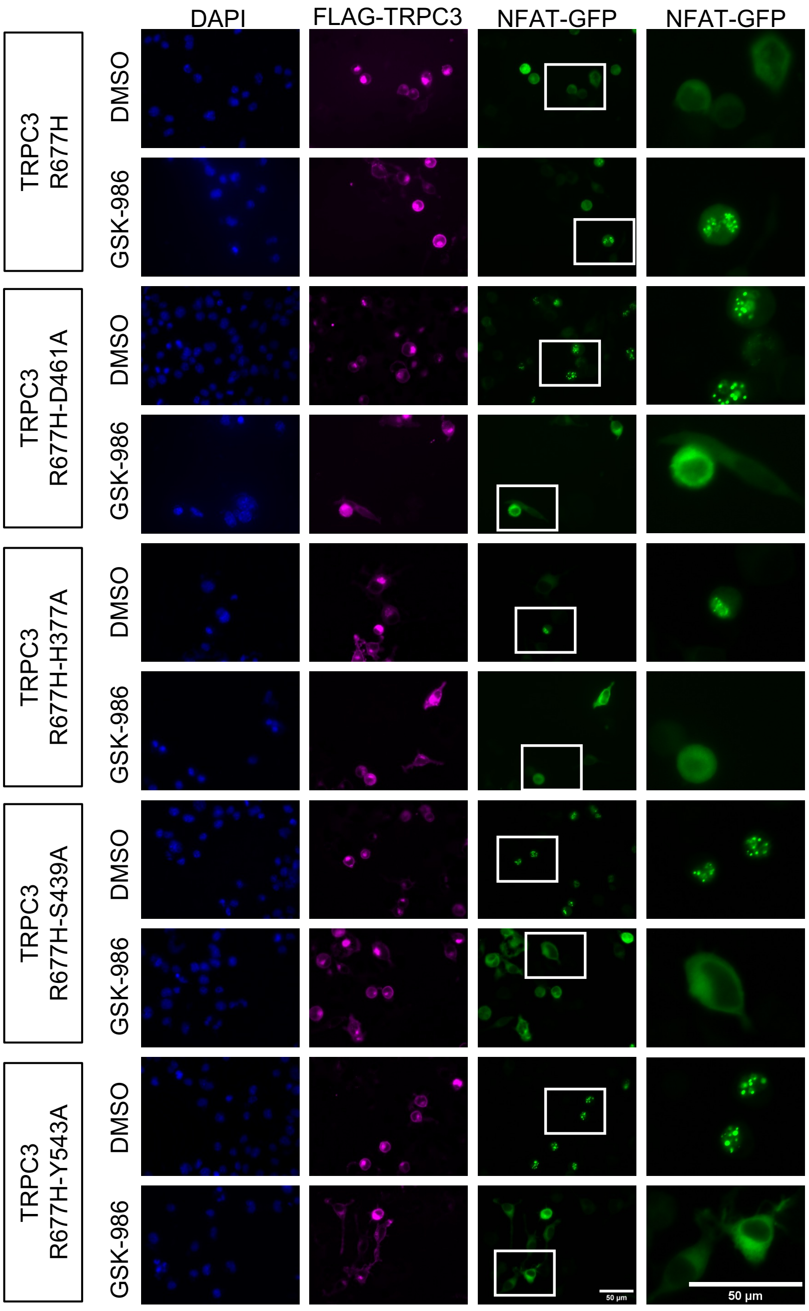
DAPI
FLAG-TRPC3
NFAT-GFP
NFAT-GFP
TRPC3 R677H
TRPC3 R677H-D461A
TRPC3 R677H-H377A
TRPC3 R677H-S439A
TRPC3 R677H-Y543A
DMSO
GSK-986
50 µm

**E**

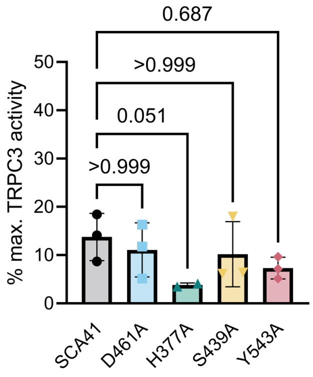
% max. TRPC3 activity
0
10
20
30
40
50
0.687
>0.999
0.051
>0.999
SCA41
D461A
H377A
S439A
Y543A

**Figure S4. GSK-986 is stable in the S1-S3 pocket in simulation but is not impaired by mutations to predicted interacting residues**. **(A)** Top 3 scoring QuickVina-W GSK-986 poses in the S1-S3, SAR7334 binding pocket. **(B)** Initial NPT GSK-986 poses compared with output poses from the first simulation run for the top 3 scoring poses: Pose 1, 2, and 3. Potential interacting residues are labelled for the initial NPT and post-simulation (Run 1) position for each pose. In total, 3 x 20-ns simulations were run for each of the three poses. Residues were identified as potential interactors for Pose 1 and 3, respectively; Pose 2 did not appear to make specific contacts with any residue. **(C)** Root-mean-squared deviation analysis for 3 x 20-ns simulations for Pose 1, Pose 2, and Pose 3. **(D)** Representative 40x magnification NFAT images of treatment of the S1-S3 binding site mutants on the SCA41 TRPC3 background with 1 μM GSK-986 or DMSO as control. Image windows are expanded to show NFAT localisation in individual cells. Background subtraction was performed equally across all images using ImageJ. Scale bars are 50 μm. **(E)** None of the tested mutants impaired inhibition of SCA41 by GSK-986 in the cell-based NFAT assay. Kruskal-Wallis test followed by Dunn's multiple comparisons test against TRPC3 R677H as control. m=2-3 independent biological replicates, each being the percentage of cells with nuclear-located GFP-NFAT across 50-200 cells. Scatter points represent biological replicates and bar graph shows mean ± standard deviation (SD). Exact p-values shown in figure.
